# Sex differences in behavioral and brainstem transcriptomic neuroadaptations following neonatal opioid exposure in outbred mice

**DOI:** 10.1101/2021.04.02.438265

**Authors:** Kristyn N. Borrelli, Emily J. Yao, Will W. Yen, Qiu T. Ruan, Melanie M. Chen, Julia C. Kelliher, Carly R. Langan, Julia L. Scotellaro, Richard K. Babbs, Jacob C. Beierle, Ryan W. Logan, William Evan Johnson, Elisha M. Wachman, Alberto Cruz-Martín, Camron D. Bryant

## Abstract

The opioid epidemic led to an increase in the number of Neonatal Opioid Withdrawal Syndrome (**NOWS**) cases in infants born to opioid-dependent mothers. Hallmark features of NOWS include weight loss, severe irritability, respiratory problems, and sleep fragmentation. Mouse models provide an opportunity to identify brain mechanisms that contribute to NOWS. Neonatal outbred Swiss Webster Cartworth Farms White (CFW) mice were administered morphine (15mg/kg, s.c.) twice daily for postnatal days (P) 1-14, an approximate of the third trimester of human gestation. Male and female mice underwent behavioral testing on P7 and P14 to determine the impact of opioid exposure on anxiety and pain sensitivity. Ultrasonic vocalizations (USVs) and daily body weights were also recorded. Brainstems containing pons and medulla were collected during morphine withdrawal on P14 for RNA-sequencing. Morphine induced weight loss from P2-14, which persisted during adolescence (P21) and adulthood (P50). USVs markedly increased at P7 in females, emerging earlier than males. On P7 and P14, both morphine exposed female and male mice displayed hyperalgesia on the hot plate and tail flick assays, with females having greater hyperalgesia than males. Morphine-exposed mice exhibited increased anxiety-like behavior in the open-field arena at P21. Transcriptome analysis of the brainstem (medulla plus pons), an area implicated in opioid withdrawal and NOWS, identified pathways enriched for noradrenergic signaling in females and males. We also found sex-specific pathways related to mitochondrial function and neurodevelopment in females and circadian entrainment in males. Sex-specific transcriptomic neuroadaptations implicate unique neurobiological mechanisms underlying NOWS-like behaviors.

**SIGNIFICANCE STATEMENT:** Neonatal opioid withdrawal syndrome (NOWS) is a poorly understood condition that has both a genetic and environmental component and is thought to be mechanistically distinct from opioid withdrawal in adults. The development of murine models for measuring neurobehavioral responses is critical for informing the neurobiological adaptations underlying NOWS. Using outbred mice that more closely model human genetic variation, we discovered a surprising degree of sexual dimorphism in behavioral timing and severity of NOWS-model behaviors as well as transcriptomic adaptations in brain tissue that together suggest distinct mechanisms and sex-specific therapeutics for reversing withdrawal symptoms and restoring brain function.

## INTRODUCTION

The United States continues to be amid an opioid epidemic, accompanied by a surge in babies exposed to opioids *in utero* (Jansson and Patrick, 2019) and a rise in rates of Neonatal Withdrawal Syndrome (NOWS) (Milliren et al., 2018). NOWS is characterized by low birth weight, reduced weight gain, severe irritability, sleep fragmentation and restlessness, hypertonia, elevated pain sensitivity, and high-pitched cry, among others (Sutter et al., 2014). Current standard of treatment for NOWS includes tapered dosing of opioids over days or weeks, and other interventions that promote mother-infant bonding (Ryan et al., 2019; Wachman et al., 2018). Current treatments for NOWS primarily attenuate withdrawal severity for the infant and are believed to mitigate the potential long-term impact on mood, cognition, and pain. Further understanding the impact of opioids on development is critical for the discovery, design, and implementation of new treatments and interventions for NOWS.

Several animal models for NOWS have been developed (Byrnes and Vassoler, 2018; Chen et al., 2015; Fodor et al., 2014), differing in time of exposure (*i.e.,* prenatally, postnatally, or both), specific opioid (morphine, heroin, methadone, oxycodone, and/or buprenorphine), and dosing regimen (*e.g.,* injections, pellets, mini-pumps) (Byrnes and Vassoler, 2018; Minakova et al., 2021; Robinson et al., 2019). The physiological and behavioral consequences of opioids on development depends on the timing, duration, and route of drug administration (Byrnes and Vassoler, 2018; Fodor et al., 2014). In rodents, the period between postnatal days 1 through 14 (P1-14) is an approximate of the third trimester of human gestation (Barr et al., 2011), characterized by rapid, extensive synaptogenesis and myelination in the brain (Semple et al., 2013). For example, morphine administration between P1-14 (Robinson et al., 2019) in mice induced neurodevelopmental delays, and effects on behaviors later during adolescence and adulthood, including marble burying and morphine locomotor sensitization behaviors (Robinson et al., 2019). Early neonatal opioid exposure in rodents has several key advantages: 1) opioid exposure during third trimester is associated with higher risk for NOWS in human neonates (Desai et al., 2015), providing strong translational relevance; 2) neonatal exposure during third trimester equivalent is sufficient to produce symptoms of withdrawal in mice and rats (Jones and Barr, 2001, 1995; McPhie and Barr, 2009; Robinson et al., 2019); 3) postnatal delivery of opioids enable accurate dosing to each pup; 4) avoiding maternal opioid exposure reduces the potential variance introduced by opioid-dependent alterations in maternal caretaking behaviors; and 5) the short-term and long-term effects of opioids on the brain and behavior can be assigned to a specific developmental period using discrete, timed administration of opioids.

Given these advantages, we employed a similar administration regimen as previously described (Robinson et al., 2019), to further investigate the consequences of neonatal opioid exposure in mouse model of NOWS, including ultrasonic vocalizations, anxiety and pain behaviors. To gain insight into the neurobiological adaptations related to NOWS, we also investigated the transcriptional alterations in the brainstem (medulla plus pons), a region of the brain implicated in neonatal opioid withdrawal (Jones and Barr, 2001; McPhie and Barr, 2009) (Maeda et al., 2002) and hyperalgesia (Bederson et al., 1990; Kaplan and Fields, 1991; Ossipov et al., 2005; Vanderah et al., 2001; Vera-Portocarrero et al., 2011; Xie et al., 2005). We used female and male outbred CFW mice (Swiss Webster) because of their increased genetic heterozygosity that better models humans (Tuttle et al., 2018), which can be leveraged in future studies to conduct high-resolution genetic mapping of NOWS-related phenotypes (Gonzales and Palmer, 2014; Parker et al., 2016).

## MATERIALS AND METHODS

### Mice

All experiments in mice were conducted in accordance with the NIH Guidelines for the Use of Laboratory Animals and were approved by the Institutional Animal Care and Use Committee at Boston University (AN-15607; PROTO201800421). Outbred Carworth Farms White (CFW) mice (Swiss Webster) were purchased from Charles River Laboratory at 8 weeks of age. Breeders were paired after one week of habituation to the vivarium. Each breeder mouse was from a different litter, thus minimizing relatedness within a breeder pair (Parker et al., 2016). Laboratory chow (Teklad 18% Protein Diet, Envigo, Indianapolis, IN, USA) and tap water were available *ad libitum.* Breeder cages were provided with nestlets. A maximum of three sequential litters per breeder were used in the study. Mice were maintained on a 12:12 light-dark cycle (lights on, 0630hr). Phenotyping was conducted during the light phase between 900 h and 1200 h. A power analysis was used to estimate the required sample size for 95% power at an alpha of 0.05 based on thermal nociception response on P7 described below. Based on the predicted effect size and power calculations, sample sizes were composed of a minimum of 12 mice per condition (6 mice per sex).

### Morphine treatment regimen from P1 through P14

Morphine sulfate pentahydrate (Sigma-Aldrich, St. Louis, MO USA) was dissolved in sterilized physiological saline (0.9%) in a 0.75 mg/ml morphine solution for systemic administration via subcutaneous (s.c.) injections. Pups were injected with morphine (15 mg/kg, 20 ul/g volume). From P1 to P14, injections were administered twice daily (0730hr and 1600hr). Pups were randomly assigned to either morphine or saline groups (20 ul/g, s.c.). On P7, pups underwent behavioral phenotyping, as described below, 16 h following morphine administration (Fig. 1A). Systemically administered morphine reaches a peak plasma concentration at 15 min with a half-life of ∼37 min (Boasen et al., 2009). Thus, the behavioral effects of opioids are attributed to withdrawal at 16hrs following last administration, rather than the acute effects of morphine. Twice daily administration of morphine then continued until P14 when pups underwent behavioral phenotyping, followed by morphine administration at 1000 h and 1600 h. On P15, 16hrs following final morphine administration, mice were sacrificed at 1000 h by live, rapid decapitation, and brain tissue was immediately extracted.

**Figure 1.**
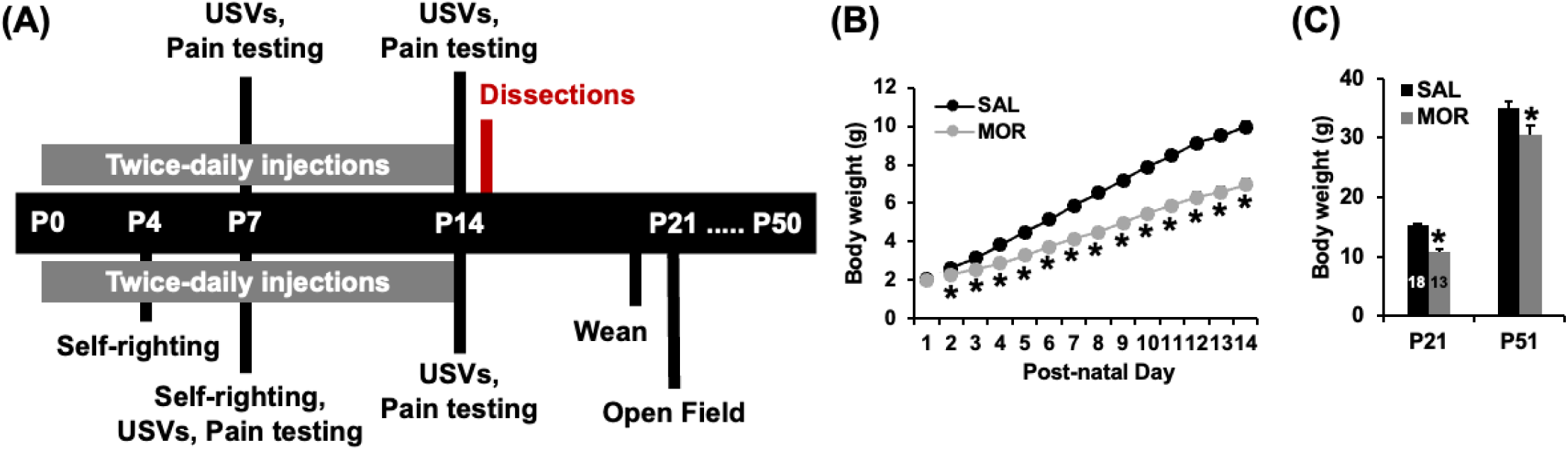
Experimental timeline and reduced weight gain following repeated neonatal morphine exposure. **(A):** Mice underwent assessment for USVs and thermal nociception. A subset of mice also underwent self-righting assessment at P4 and P7 as well as anxiety-like behavior in the open field at P21. **(B):** In examining changes in body weight, a 2-way ANOVA (treatment, sex) revealed no main effect of treatment on P1 (saline n=29, morphine n=25; F_1,51_ = 0.03, p = 0.86). Thus, there was no random effect of treatment. There was the expected main effect of sex (F_1,51_ = 12.49, p = 8.8 x 10^-4^) but no treatment x sex interaction (F_1,51_ = 0.57; p = 0.46). Body weight was subsequently analyzed from P2-P14 via mixed-model ANOVA, which revealed a main effect of treatment (saline n=41, morphine n=35; F_1,72_ = 98.28, p = 4.26 x 10^-15^), day (F_12,864_ = 1605.67, p < 2 x 10^-16^), and a treatment x day interaction (F_12,864_ = 73.27, p < 2 x 10^-16^). *Post hoc* pairwise t-tests revealed significantly lower body weight in the morphine group (**\*** *post hoc* t-tests all p < 3 x 10^-4^; α_adjusted_ = 0.0038) **(C):** In examining body weight in the subset of mice that were assessed at P21 and P51, there was a main effect of treatment (F_1,27_ = 26.51, p = 2.04 x 10^-5^), sex (F_1,27_ = 25.09, p = 2.97 x 10^-5^), Day (F_1,27_ = 1179.85, p < 2 x 10^-16^), and day x sex interaction (F_1,27_ = 30.21, p = 8.04 x 10^-6^). In support of the treatment effect, morphine mice showed a significant decrease in body weight compared to saline mice at both P21 (**\***t_1,29_ = - 7.33; p = 4.55 x 10^-8^) and at P50 (**\***t_1,29_ = -2.27; p = 0.03).

### General procedures

Experimenters were blinded to saline or morphine conditions behavioral testing on P7 and P14. Pups were habituated to the testing room for at least an hour prior to testing. Immediately before testing pups were transferred to a new cage containing a heat pad. Dams and sires remained in their home cages in the testing room while the pups were tested. Activity in sound attenuating chambers during ultrasonic vocalization recording and on the hot plate were videoed using infrared cameras (Swann Communications U.S.A. Inc., Santa Fe Springs, CA, USA) and tracked with ANY-maze tracking software (Stoelting Co., Wood Dale, IL, USA).

### Ultrasonic Vocalizations (USVs) in P7 and P14 mice

During maternal separation, withdrawal from repeated morphine administration increases USVs in rat pups (Barr and Wang, 1992). Prior to injections on P7 and P14, each pup was placed into a Plexiglas box (43 cm L x 20 cm W x 45 cm H; Lafayette Instruments, Lafayette, IN, USA) placed within sound-attenuating chambers (Med Associates, St. Albans, VT, USA). USVs were recorded using Ultrasound Recording Interface (Avisoft Bioacoustics UltrasoundGate 816H, Glienicke, Germany) for 10min. After 10 min, pups were removed from the boxes, weighed, and placed in a holding cage with a heating pad prior to pain testing.

We used an unsupervised approach to categorize and compare syllable repertoires based on the similarity of their spectrotemporal patterns (Mouse Ultrasonic Profile Extraction; MUPET (Van Segbroeck et al., 2017)). Briefly, MUPET determines specific syllable types present in mouse USVs by analyzing their entire frequency structure, independent of syllable fundamental frequency or amplitude. Extracted syllable shapes were centered along time and frequency and represented low dimension representations of USV patterns that captured their spectral shape and maintained the key spectro-temporal features (Bertrand et al., 2008; Joder and Schuller, 2012). Mean syllable duration was determined empirically from the distribution of syllable duration values in the saline group, then used for unsupervised analyses. Syllable thresholds were set at a minimum of 8ms and maximum of 200 ms. Optimal syllable repertoire size was determined using multiple measures from MUPET (Bayesian information criterion (BIC), average log likelihood, overall repertoire modeling score and goodness-of-fit (data not shown)). The diagonal of the Cross Repertoire Similarity Matrix which provides the correlations between syllable unit pairs from two different repertoires was used to compare the shapes of syllable units between experimental groups.

### Thermal nociception in P7 and P14 mice

Upon completion of USV recordings, the first pup was removed from the holding cage and placed in a Plexiglas cylinder (15 cm diameter; 33 cm H) on a 52.5°C hot plate (IITC Life Science INC., Woodland Hills, CA, USA). On P7, the latency for the pup to turn onto their back was used as an indication of nociceptive detection. The nociceptive response on P14 was the latency to flick or lick the forepaw or hindpaw, or jump. On P14, majority of pups displayed a hindpaw lick. Pups were moved immediately following a pain response or following 30s with no response. We also calculated the velocity of travel of pups on the hot plate prior to the presentation of a nociceptive response by dividing the distance travelled (ANY-maze software) by the nociceptive latency. Immediately following the hot plate, each pup was gently scruffed and the distal half of its tail was quickly lowered into a 48.5° hot water bath (LX Immersion Circulator, PolyScience, Niles, IL) and the tail flick latency was recorded. For hot plate velocity, we excluded a total of four pups from the analysis (n = 2 saline, n = 2 morphine) due to errors with the tracking software (n=39 saline and n=33 morphine).

### Tattooing in P7 mice

Following thermal nociception testing on P7, pups were tattooed for identification (ATS-3 General Rodent Tattoo System, AIMS™, Hornell, NY, USA). Pups were then injected with either morphine or saline then returned to their home cage with the dam and sire.

### Self-righting in P4 and P7 mice

A subset of pups was also tested for self-righting as a developmental milestone (Robinson et al., 2019). On P4 and P7, each mouse was placed onto their back with all paws facing upward, and gently stabilized by a finger. Latency for the mouse to right itself and place all paws on the surface was recorded.

### Open field arena

A subset of mice underwent testing for anxiety-like behavior in the open field arena on P21. Mice could roam freely in Plexiglas boxes (20cm W x 43cm L x 46cm H) for 5min. Behavior was video recorded then later scored using ANY-maze software.

### Statistical analysis

Data analysis was performed in R (https://www.r-project.org/). Except for open field measures, all phenotypes were analyzed using mixed-model ANOVAs, with condition and sex as independent factors and either age or day as categorical repeated measures. Distance and percent time in center of open field arena were analyzed using 2-way ANOVA, with treatment and sex as independent variables. *Post hoc* unpaired Student’s t-tests were conducted as indicated. A Bonferroni correction was used for USVs/min, USV distance/min, and P21 open field distance/min. On P1, body weights were analyzed using separate 2-way ANOVA due to partially missing data. Body weights from P2 through P14 analyzed using 2-way repeated measures ANOVA (treatment and sex as independent variables, age as the repeated measure).

### Brainstem RNA-sequencing (RNA-seq)

On P15, 16 h following the final morphine injection, mice were sacrificed, and brains were quickly removed to collect the brainstem containing the medulla and pons. Brainstems were transferred to tubes containing RNA-later, stored at 4°C, then several days later, lightly blotted and transferred to new tubes to store at -80°C until RNA extraction. RNA was extracted using Trizol (Qiagen), ethanol precipitation, filtering columns (Qiagen), and elution with sterile, double deionized water (Yazdani et al., 2015). Samples were diluted to 50 ng/ul. RNA library preparation (poly-A selection) and RNA-seq was conducted at the University of Chicago Genomics Facility on an Illumina NovaSEQ6000 using a NovaSEQ SP-100 bp flowcell/reagent cassette. The 12 multiplexed, pooled samples (6 mice from saline and 6 mice from morphine groups) were sequenced (100 bp single-end) on a single lane of the two-lane flowcell. We used the R/Bioconductor package “scruff” for data preprocessing, including demultiplexing, read alignment, read counting, quality checking and data visualization (Wang et al., 2019). Reads were trimmed for quality using Trimmomatic (Bolger et al., 2014). Trimmed reads were then aligned to the mm10 mouse reference genome (Ensembl) to generate BAM files for alignment using STAR (Dobin et al., 2013). The featureCounts read summarization program was used to count reads mapping to the “exon” feature in a GTF file obtained from Ensembl (GRCm38). Differential gene expression analysis of normalized read counts from Rsubread (Liao et al., 2019) was performed in R using the exactTest function in edgeR (Robinson et al., 2010). RIN scores from the Bioanalyzer ranged from 8.2 to 8.9, indicating high quality RNA for sequencing. We obtained an average of 43.7 million reads per sample (range: 35-60 million reads per sample). 97.7% to 98.6% of these reads had a perfect barcode match. Phred scores were all 36.4. Greater than 90% of the reads mapped uniquely to a single locus. FASTQ files and normalized read counts are available for download in NCBI Gene Expression Omnibus (GSE141066).

### Gene network and pathway enrichment

Network analysis and plot generation were performed using Cytoscape software. Gene expression data sets (containing log fold-change (LogFC) and p-values) were uploaded for pathway analysis and networks were plotted using known gene interactions imported from the STRING database. All genes with unadjusted p < 0.10 were uploaded for network analysis, which enabled the detection of more broad patterns of gene expression within specific functional networks. We first analyzed the effect of morphine on gene expression within the combined analysis which used sex as a covariate. For the combined analysis, we plotted the top six GO Biological Process networks that contained fewer than 50 genes to simplify viewing of the network plot and avoid overly broad enrichment terms. We then probed sex-specific effects of morphine by separately analyzing males and females. For males and females, we determined the top KEGG enrichment terms and identified the gene-gene interactions as determined by STRING.

### Alternative splicing

Analysis was performed using the R package ASpli (Mancini et al., 2020) (Bioconductor version 1.14.0). We first performed feature extraction of the GTF annotation files to retrieve information regarding gene structure, including intron and exon bins, junctions, and intron flanking regions. Using the BAM files, we mapped RNA-seq reads to the annotated genome to determine read count density for each extracted feature. We determined differential intron/exon bin usage to identify alternative splicing events. We assessed the main effect of MOR treatment on splicing events in sex-collapsed, female-only, and male-only datasets. We also identified splicing events influenced by interactive effects of condition and sex.

### Correlation of read counts with P14 behavioral phenotypes

To gain further insight into the molecular adaptations contributing to the behavioral adaptations induced by morphine, we examined the correlation between gene expression levels and phenotypes collected on P14. We specifically sought to identify genes that were differentially expressed in the morphine condition collapsed across sex and genes with normalized read counts that significantly correlated with P14 phenotypes (hot plate latency and velocity, tail withdrawal latency, USVs/min, distance traveled during USV recordings, and body weights. Pearson product moment correlation coefficients were calculated between the gene expression data and each of these phenotypes.

## RESULTS

### Reduced weight gain following neonatal morphine exposure

Hallmarks of NOWS in human neonates are low birth weight and reduced weight gain. On P1, prior to morphine administration, body weights were similar across mice (Fig. 1B; n=54). By P2, pups that were administered morphine (n=35) had significantly lower body weights relative to saline pups (n=41), persisting through P14 (Fig. 1B). Lower body weights were also evident in morphine-exposed pups (n=18) compared to the saline group (n=18) at P21 and P51 in a subset of mice tested for self-righting and open field behavior (Fig. 1C). Body weights for pups that died prior to P14 (6 of 41 morphine exposed pups, ∼15%) were excluded from further analyses. Pups were identified as female or male on P14, so sex-specific mortality could not be determined. Thus, morphine administration from P1-14 lead to acute reductions in body weights and marked lower weight gain that persisted into adulthood.

### Altered self-righting latency in a sex-specific and development-dependent manner following repeated neonatal morphine exposure

Self-righting from back to abdomen is used to test milestones associated with motor development. On P4, pups administered morphine displayed significantly prolonged latencies to self-right compared to saline mice (Fig. 2). By P7, there was a significant decrease in self-righting latency in males administered morphine compared to P4 (Fig. 2). However, females administered morphine still exhibited prolonged self-righting latencies compared to controls and males (Fig. 2). Together, our results suggest a morphine-induced developmental delay that was more persistent in females exposed to morphine compared to males.

**Figure 2.**
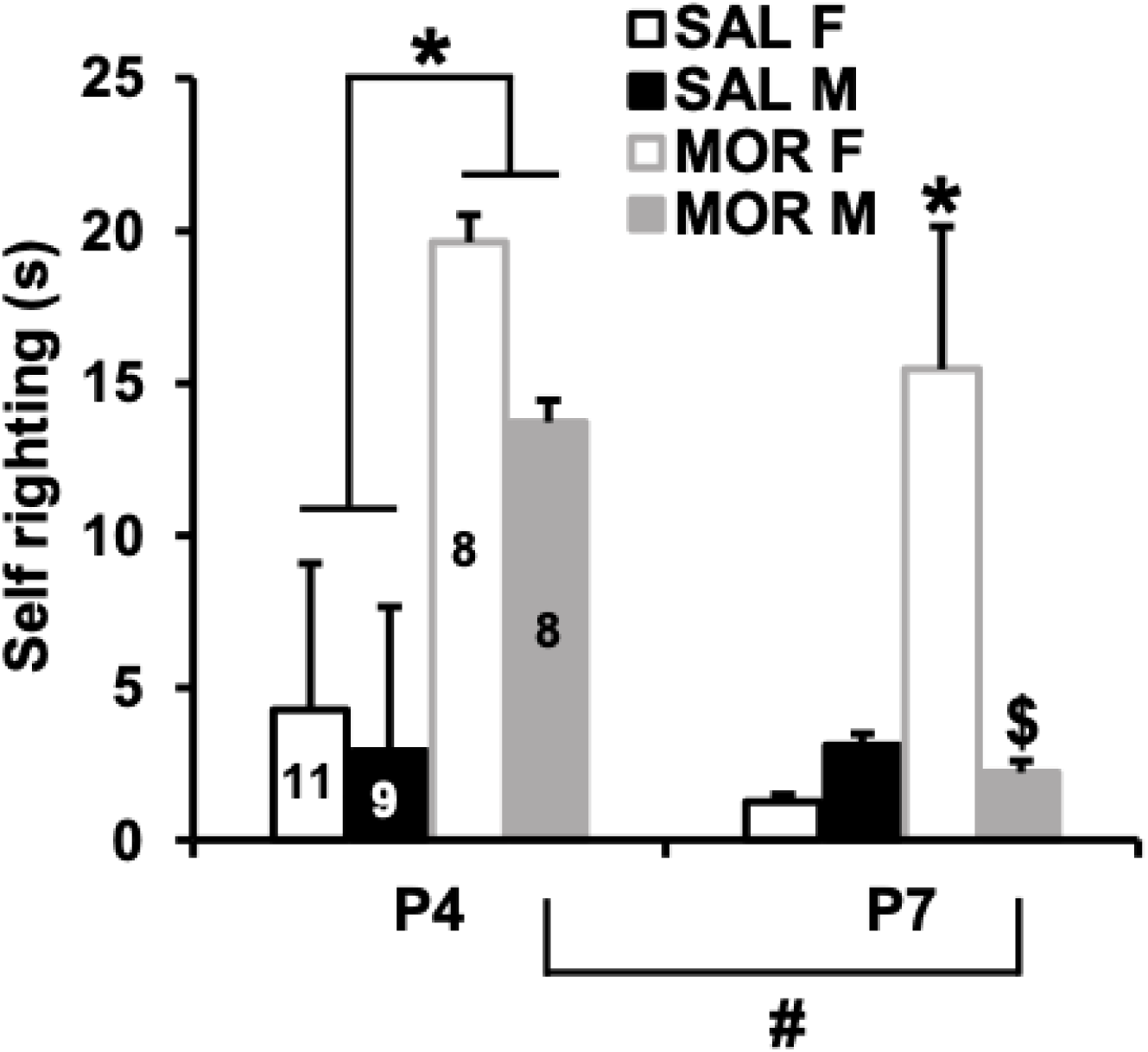
Altered self-righting latency in a sex-specific and development-dependent manner following repeated neonatal morphine exposure. In examining the latency to self-right on P4 and P7, there was a main effect of postnatal day (F_1,32_ = 5.62, p = 0.024), treatment (F_1,32_ = 24.81; p = 2.1 x 10^-5^), sex (F_1,32_ = 4.38, p = 0.044), a treatment x sex interaction (F_1,32_ = 6.11, p = 0.019), but no significant treatment x sex x day interaction (F_1,32_ = 2.02; p = 0.17). On P4, there was a main effect of treatment (F_1,32_ = 18.13, p = 1.7 x 10^-4^). Morphine exposed pups showed a significantly longer latency to self-right than saline exposed pups (t_1,34_ = 4.27, **\***p = 1.48 x 10^-4^). On P7, there was a main effect of Treatment (F_1,32_ = 8.47, p = 0.007), Sex (F_1,32_ = 4.48, p = 0.04), and a treatment x sex interaction (F_1,32_ = 10.55, p = 0.003) that was explained by morphine exposed females showing a significantly longer latency to self-right compared to their saline female counterparts (**\***t_1,17_ = 3.58, p = 0.0023) and compared to morphine exposed males (**$**t_1,14_ = 2.81; p = 0.01). Morphine exposed males did not differ significantly from saline males (t_1,15_ = 0.42; p = 0.68). Finally, whereas morphine exposed males showed a significant improvement (decrease) in the latency to self-right between P4 and P7 (t_1,14_ = 2.39; p = 0.03), morphine exposed females did not (t_1,14_ = 0.63; p = 0.54).

### Increases in ultrasonic vocalizations (USVs) during maternal separation following repeated neonatal morphine exposure

In rodents, USVs are associated with negative affective state and used as an indicator of opioid withdrawal (Barr et al., 2011). We found complex relationships between condition, sex, age and day in USVs. As expected, mice displayed an overall decrease in USVs from P7 through P14 (Noirot, 1966) (Fig. 3A), mainly due a greater reduction in USVs in saline mice (P14 vs. P7). Notably, morphine exposed pups displayed sustained elevations in USVs relative to saline pups (Fig. 3A). We then parsed USVs by sex and time during the recording session (10mins) on P7. During the initial 5mins of the recording session, morphine exposed females had significantly more USVs per minute than saline females (Fig. 3B). There were no differences in USVs during the initial 5mins of the recording session across groups in male pups. On P14, USVs per minute were similar between males and females administered morphine compared to the saline group (Fig. 3C). Our findings indicate morphine administration from P1-P14 leads to an overall increase in USVs with male pups displaying augmented vocalizations at P7, consistent with previous reports (Robinson et al., 2019).

**Figure 3.**
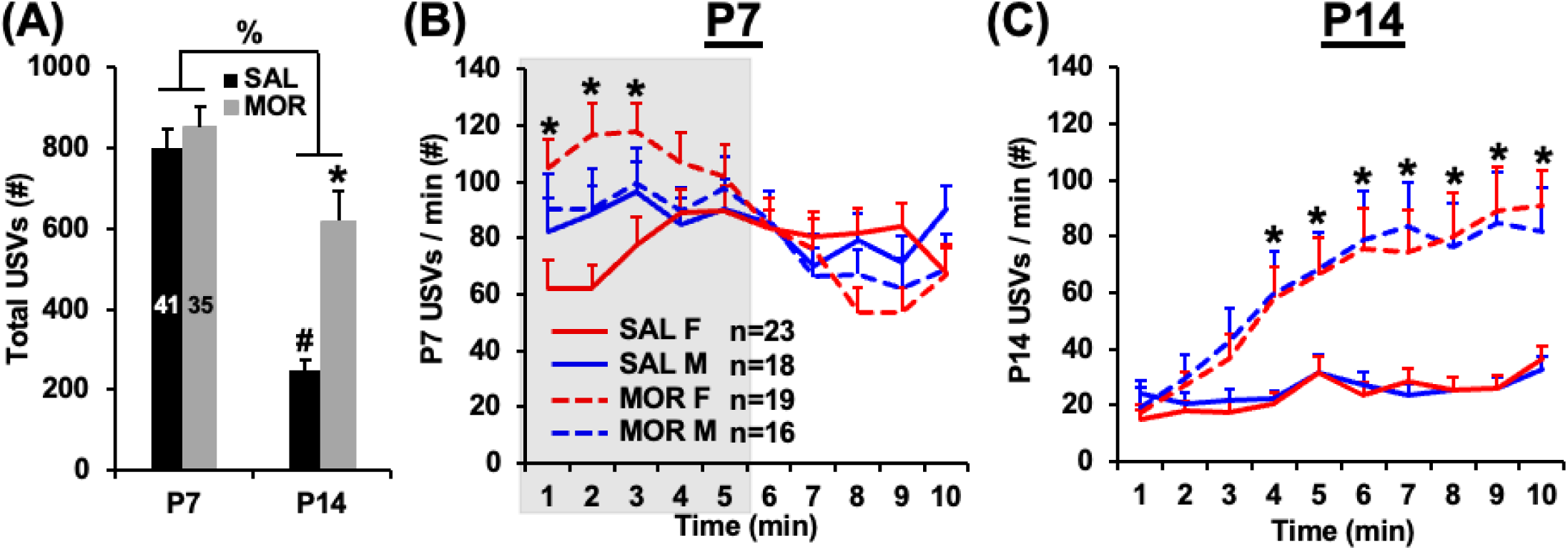
Pronounced increases in ultrasonic vocalizations (USVs) during maternal separation following repeated neonatal morphine exposure. **(A)** In examining total USVs emitted over the 10 min, a mixed-model ANOVA revealed a main effect of postnatal day (**%**F_1,71_ = 69.88, p = 3.6 x 10^-12^) and a Treatment x postnatal day interaction (F_1,71_ = 8.23, p = 0.005). There was no difference between saline and morphine exposed groups on P7 (t_74_ = 0.78, p = 0.44). Furthermore, as expected, saline exposed mice showed a reduction in the number of USVs from P7 to P14 (**#**t_1,40_ = 12.10; p = 5.96 x 10^-15^). In contrast, for morphine exposed mice, there was no significant decrease in USVs from P7 to P14. Furthermore, on P14, morphine exposed mice showed more USVs than saline mice (**\***t_74_ = 4.97; p = 4.18 x 10^-6^). **(B)** In examining USVs per min, visual inspection of the first five min indicated potential sex-specific effects. In analyzing the first five min, there was a main effect of treatment (F_1,72_ = 6.11; p = 0.016) and a nearly significant effect of time (F_4,288_ = 2.25; p = 0.064), a trending treatment x sex interaction (F_1,72_ = 2.83; p = 0.097), and a nonsignificant treatment x sex x time interaction (F_4,288_ = 1.89; p = 0.11). Female morphine mice showed a significant increase in USVs compared to female saline mice during the first three min (p = 0.0042, 3.3 x 10^-4^, and 0.0093, respectively) that survived the Bonferroni-adjusted p-value for significance (0.05/5 = 0.01_adjusted_). In contrast, there was no significant difference between male treatment groups at min 1, min 2, or min 3 (p = 0.63, 0.91, 0.85). **(C)** In examining USVs on P14 across all 10 min, there was a main effect of treatment (F_1,72_ = 17.32, p = 8.64 x 10^-5^), and time (F_9,648_ = 106.83, p < 2 x 10^-16^) and a treatment x time interaction (F_9,648_ = 24.41, p < 2 x 10^-16^). Because there was no effect of sex or treatment x sex interaction at any time point (p > 0.43), we collapsed the data across sex for each treatment and ran unpaired t-tests between the morphine exposed group and the saline group. For min 4 through min 10, the morphine exposed pups emitted significantly more USVs/min than the saline pups (*p ≤ 4.9 x 10^-4^; α_adjusted_ = 0.005).

### Altered USV syllable repertoires following repeated neonatal opioid exposure

Mouse vocalizations are characterized by diverse patterns of syllable repertoires with distinct spectrotemporal patterns. These repertoires are developmentally regulated and elicit a variety of behavioral responses from the dam that are crucial for bonding and survival, including feeding, pup retrieval, licking, and grooming. We used MUPET (Van Segbroeck et al., 2017) to determine whether morphine administration during early postnatal development caused changes in the spectrotemporal patterns of USV syllables. Morphine administration increased syllable duration, maximum frequency, and frequency bandwidth on P7 selectively in female mice (Fig. 4), suggesting sex-specific differences in developmental sensitivity to morphine. By P14, USV characteristics in syllable duration, minimum frequency, maximum frequency, and mean frequency were significantly increased in both males and females (Fig. 4). We next categorized syllable repertoires in female and male mice administered saline at P7 (Fig. 5), then arranged repertoires by sex based on spectrotemporal similarity (Fig. 5A,B). Syllable units between female and male mice were highly similar at P7 (Fig. 6C; Pearson correlation 0.90 ± 0.02).

**Figure 4.**
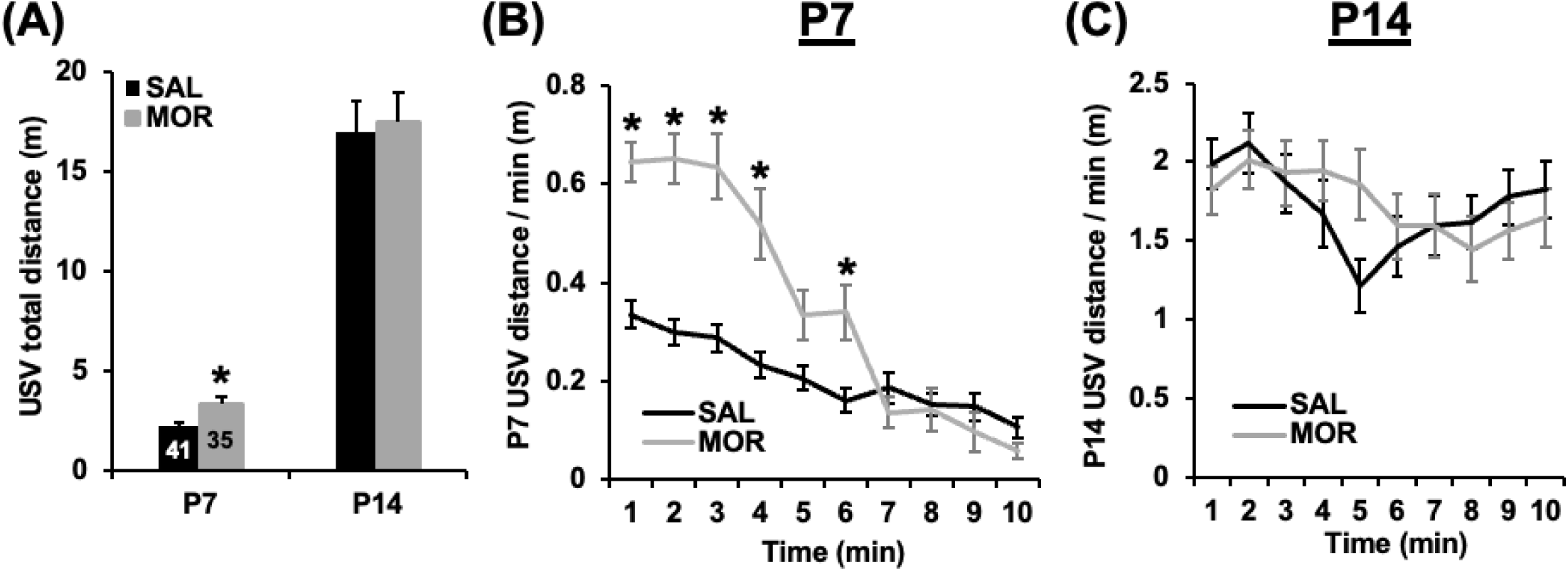
USV syllable repertoires following repeated neonatal morphine exposure. Syllable characteristics obtained from unsupervised approach to categorize syllable repertoires (MUPET). **(A)** Chronic morphine exposure led to both an increase in syllable duration in P7 female mice and P14 mice of both sexes relative to saline controls. **(B-C)** Decrease in inter-syllable interval and increase in minimum frequency in P14 mice of both sexes with morphine exposure. **(D)** Chronic morphine exposure led to both an increase in maximum frequency in P7 female mice and P14 mice of both sexes relative to saline controls. **(E)** Increase in mean frequency in morphine treated P14 mice of both sexes relative to saline controls. **(F)** Chronic morphine exposure led to both an increase in frequency bandwidth in P7 and P14 mice of both sexes relative to saline controls. We saw a significant effect of group (saline vs. morphine treatment) for Syllable duration (\**p*<0.0001), Inter-syllable interval (\*\*\*\**p*<0.0001), Maximum frequency (\*\*\*\**p*<0.0001), Mean frequency (\*\*\*\**p*<0.0001) and Frequency bandwidth (\*\*\**p*=0.0008) and a significant effect of Development x Group factors for Syllable duration (\**p*=0.0288), Inter-syllable interval (\*\*\*\**p*<0.0001), Minimum frequency (\*\*\*\**p*<0.0001), Maximum frequency (\*\**p*=0.0025) and Mean frequency (\*\*\*\**p*<0.0001). We followed Repeated Measures testing with Tukey’s multiple comparison post-test (asterisks shown in graphs). Solid pink square, Saline female group, Solid blue circle, Saline male group, Hollow pink square, Morphine female group, Hollow blue square, Morphine male group. Data as Mean ± SEM.

**Figure 5.**
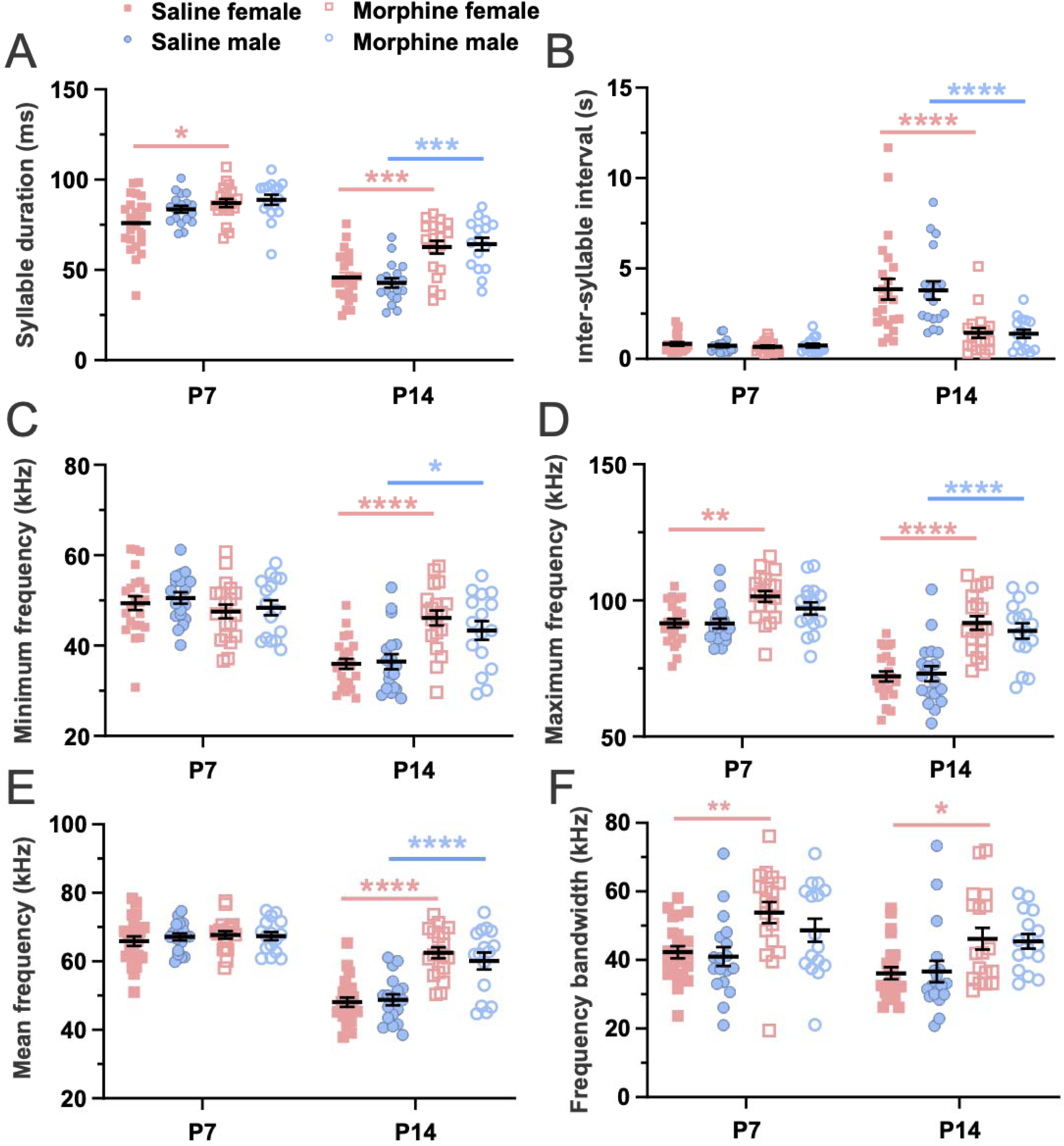
USV spectrotemporal structure was developmentally regulated. **(A-B)** Repertoires showing the 40 syllable types (average shape; top black numbers, syllable number) learned from processing recordings across the saline P7 data sets. Syllable types are listed in order of frequency of use from left to right (1-40), with the total number of syllables that are present in each syllable unit given in blue. Black number in parenthesis, original syllable number from datasets before doing comparison analysis. Red and blue squares are examples of highly similar or distinct syllables across groups, respectively. Pearson similarly scores are < 0.70 after syllable labeled with bold black square. **(C-D)** Examples of highly similar (red squares in **(A)** and **(B)**) and distinct syllables across saline P7 groups. **(E-F)** Same as in **(A)** and **(B)** but syllables learned from saline P7 and P14 data sets. **(G-D)** Examples of highly similar (red squares in **(A)** and **(B)**) and distinct syllables across saline P7 groups. **(A-H)**, the lower side of each syllable frame is 150 ms. **(I)** Similarity Matrix of the spectral types of pairs of syllables units learned from comparisons between reference group and saline P7 male mice (left panel) or saline P14 female mice (right panel). The matrix diagonal gives the Pearson correlation for sequential pairs of syllables from compared datasets ranked from most to least similar (e.g., syllable 1 in both repertories are highly similar). Bold black square in Similarity Matrix indicates highly distinct syllable units (Pearson score ≤ 0.70) and increases in saline P7/P14 comparison (left panel) relative to P7 comparisons (right panel), suggesting developmental regulation of USV spectrotemporal structure. **(J)** Mean Pearson correlation values obtained from the diagonal in comparisons of saline P7 male **(I)** and P14 groups of both sexes (**(I)** and Fig. 3C) to saline P7 female mice, “Reference Group”. Red and blue values, Pearson correlations for highly similar and distinct pairs of syllables, respectively. One-Way ANOVA followed by Tukey’s multiple comparison post-test (asterisks shown in graphs). Data as Mean ± SEM.

**Figure 6.**
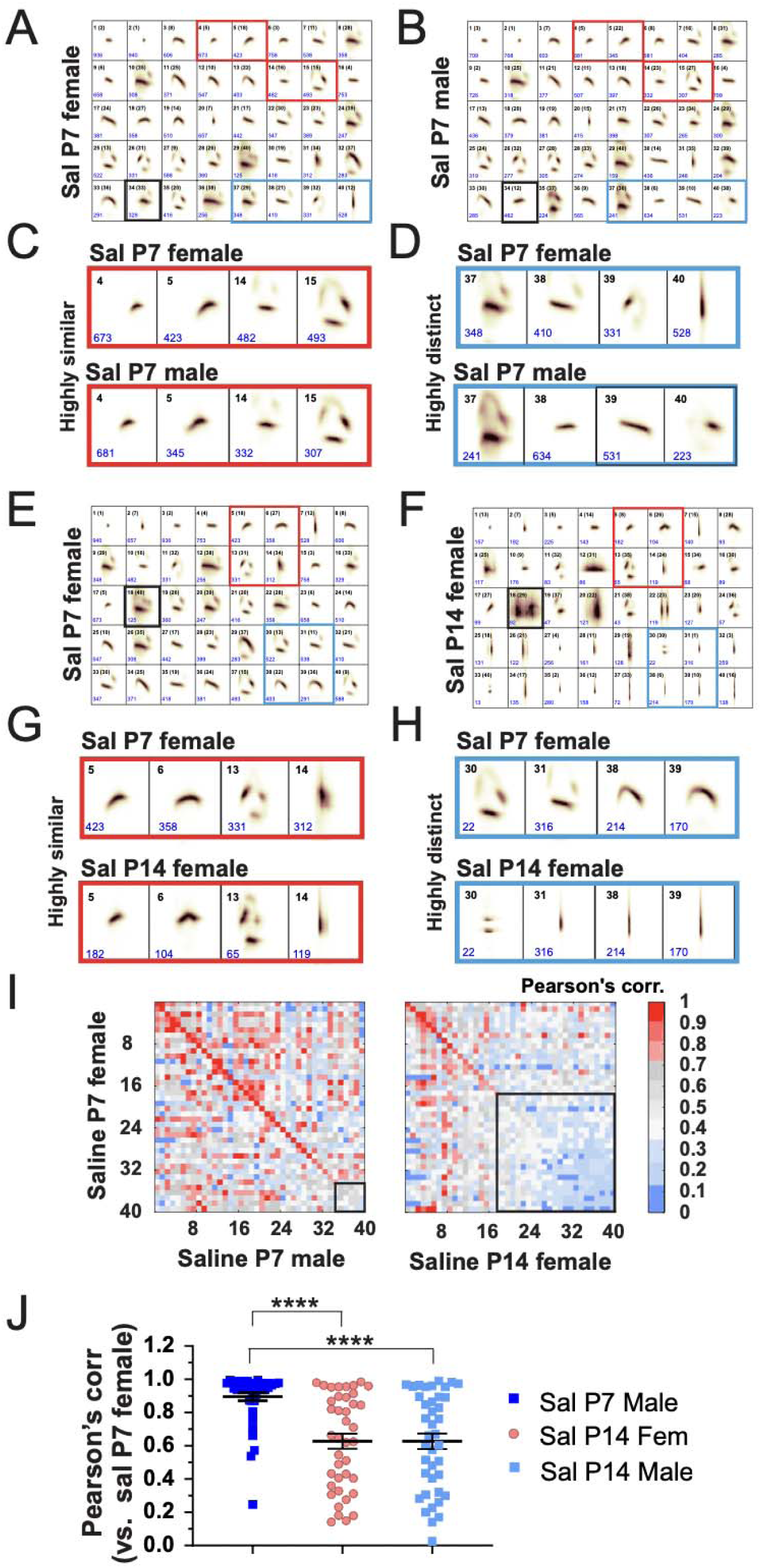
Alterations in the spectrotemporal patterns of USV syllables during the second week of postnatal development. **(A-E)** Similarity Matrices of the spectral types of pairs of syllables units learned from reference group comparison to morphine P7 female **(A)** and male **(B)**, saline P14 male **(C)**, morphine P14 female **(D)** and male **(E)**. (**G**) Mean Pearson correlation values obtained from the diagonal in comparisons of all groups to Reference Group. One-Way ANOVA followed by Tukey’s multiple comparison post hoc test (asterisks shown in graphs). Data as Mean ± SEM.

Pearson correlation values between syllable repertoires displayed a larger decrease between saline females at P7 and both saline male and female groups at P14 than to saline males at P7 (Fig. 5I,J; values obtained from the diagonal; black square, highly distinct syllable units, Pearson score ≤ 0.70). Overall, our findings suggest our approach had the sensitivity to detect developmental changes in the spectrotemporal patterns of USV syllable repertoires of early postnatal mice. In addition, these results suggest a transition in the spectrotemporal structure of USV syllables during the first two weeks of postnatal development.

Although repeated morphine administration affected USV syllables of females during the first week of postnatal development (Fig. 4), morphine had minimal effects on the Pearson correlation values obtained from the similarity matrices of either females or males relative to P7 saline groups (Fig. 5I,J; Fig. 6A,B,G). Overall, these results suggest that repeated morphine administration had no effects on the spectrotemporal structure of USVs of P7 mice of either sex. Further, Pearson correlations obtained from comparing saline females at P7 to morphine exposed mice at P14 were smaller (Fig. 6D-G) than when compared to the saline group at P14 (Fig. 5E-J). Moreover, Pearson correlations from comparisons between saline females at P7 and the morphine group at P14 (Fig. 6D-G) were not significantly different from those generated from either comparing the reference group to their P7 SAL male counterparts (Fig. 5I,J) or to saline P14 of both sexes (Fig. 5I,J; Fig. 6C,G). Therefore, early developmental morphine exposure slowed the developmental trajectory of the USV spectrotemporal structure to more closely resemble the earlier P7 repertoire. Alternatively, morphine withdrawal at P14 could mimic the apparent developmental delay of USVs. We also measured locomotor activity during USV recording sessions. On P7, mice that were previously administered morphine exhibited a marked increase in locomotory activity, an effect that was absent at P14 (Fig. 8A). Mice that were previously administered morphine displayed increased locomotor velocity (distance by min) during the initial 7 minutes of the session (Fig. 8B). There were no discernible differences in locomotor velocity between groups on P14 (Fig. 8C). Overall, locomotor velocity during USV recordings paralleled changes in USV frequency in pups that were previously administered morphine.

**Figure 7.**
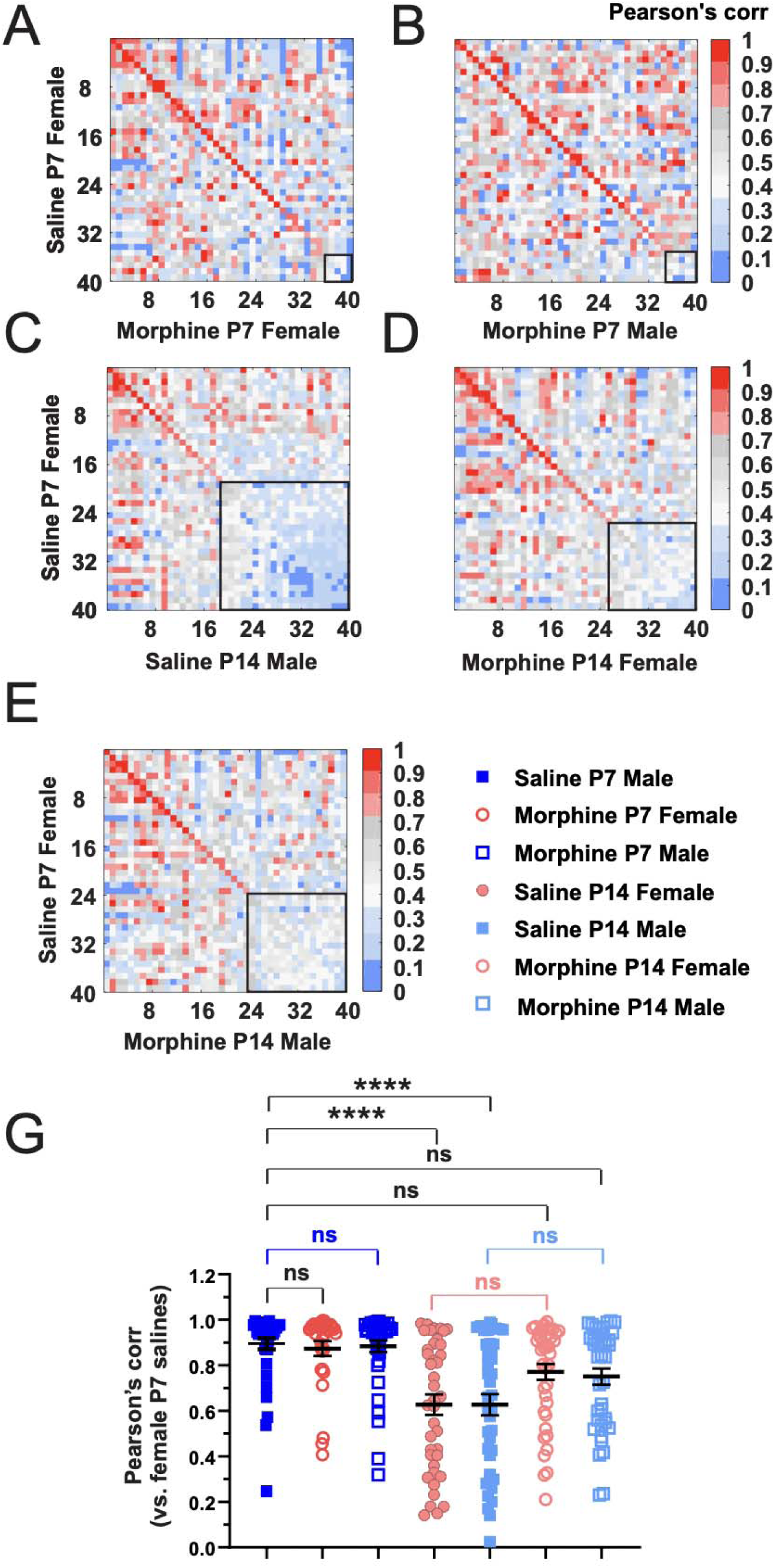
Concomitant locomotor activity during USV emissions following repeated neonatal morphine exposure. **(A)** In examining distance traveled in the USV box over 10 min, mixed-model ANOVA (Treatment and sex as factors, day as a repeated measure) revealed a main effect of postnatal day (F_1,72_ = 197.90, p < 2 x 10^-16^), but no effect of treatment, sex, and no interaction (p’s≥0.31). Morphine exposed pups traveled a significantly greater distance than saline exposed pups at P7 (t_74_ = 3.05, p = 0.003) but not at P14 (t_1,74_= 0.24, p = 0.81). **(B)** In examining the time course of the increase in USV distance in morphine-treated pups at P7, a mixed-model ANOVA revealed a main effect of treatment (F_1,72_ = 15.59, p = 1.81 x 10^-4^), time (F_9,648_ = 53.43, p < 2 x 10^-16^), and a treatment x time interaction (F_9,648_ = 17.59, p < 2 x 10^-16^). During min 1 through 4 and min 6, the morphine group traveled significantly more distance per minute than the saline group (*unpaired t-tests, all p < 0.002; α_adjusted_ = 0.005). **(C)** In examining distance traveled on P14, there was a main effect of time (F_9,639_ = 4.68, p = 4.93 x 10^-6^) and a treatment x time interaction (F = 2.49, p = 0.008). However, in contrast to P7, there was no significant difference between the saline and morphine groups at any time point (p > 0.03, α_ adjusted_ = 0.005).

**Figure 8.**
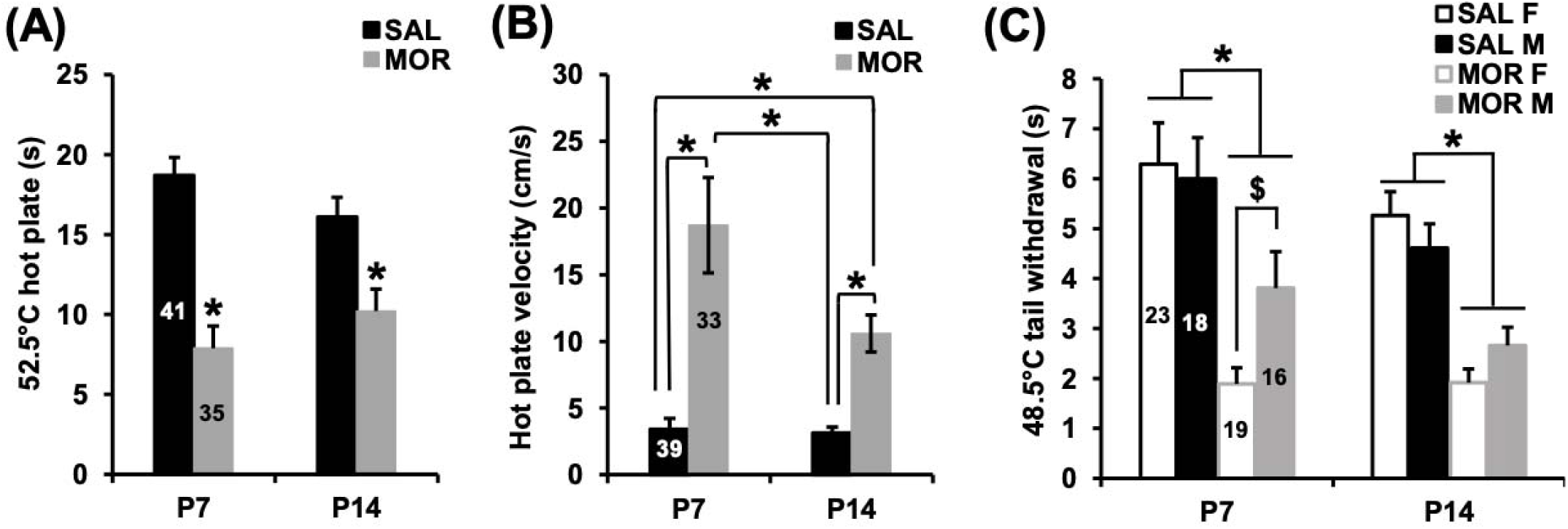
Thermal hyperalgesia following repeated neonatal morphine exposure. **(A)** In examining thermal nociceptive sensitivity on the 52.5°C hot plate assay on P7, a mixed-model ANOVA (treatment and sex as independent variables; postnatal day as a repeated measure) revealed a main effect of treatment (F_1,72_ = 51.00, p = 6.07 x 10^-10^) but no effect of sex (F_1,72_ = 0.62; p = 0.43), postnatal day (F_1,72_ = 0.06; p = 0.80), or any interactions (p≥ 0.08). Morphine exposed pups showed hyperalgesia as indicated by a significantly shorter latency to respond on both P7 (t_1,74_ = 6.13, p = 3.98 x 10^-8^) and P14 (t_1,74_ = 3.21, p = 0.002). **(B)** In examining thermal nociceptive sensitivity with the 48.5°C tail withdrawal assay, mixed-model ANOVA (treatment and sex as factors, day as a repeated measure) indicated a main effect of treatment (F_1,72_ = 58.33, p = 7.28 x 10^-11^), postnatal day (F_1,72_ = 4.01, p = 0.05), and a treatment x sex interaction (F_1,72_ = 4.99, p = 0.03). On both P7 and P14, morphine exposed pups (collapsed across males and females) showed a decrease in latency to tail withdrawal (P7: **\***t_74_ = 4.64, p = 1.47 x 10^-5^; P14: **\***t_74_ = 6.40, p = 1.25 x 10^-8^). Interestingly, on P7, morphine exposed females showed a more robust hyperalgesia as indicated by significantly shorter tail withdrawal latency than morphine exposed males (**$**t_1,33_ = 2.55, p = 0.02).

### NOWS led to thermal hyperalgesia

Morphine administration from P1-P14 led to hyperalgesia. At P7 and P14, morphine exposed pups showed a shorter latency to respond to a nociceptive stimulus on the hot plate and tail withdrawal assays (Fig. 8A,C). Morphine females displayed significantly enhanced hyperalgesia (shorter latencies) compared to morphine males (Fig. 8C). For P14 tail withdrawal, although hyperalgesia was still evident overall (Fig.8C), there was no longer a significant sex difference. Morphine exposed pups also displayed greater velocities on the hot plate prior to displaying a nociceptive response (Fig. 8B).

### NOWS led to increased anxiety-like behavior in the open field at P21

We measured distance traveled and time spent in the open field arena at P21 in a subgroup of mice that were previously administered saline (n=20) or morphine (n=15). Prior morphine administration decreased distance traveled and percent time spent in the center of the open field arena compared to saline mice (Fig.9A-B), reflective of increased anxiety-like behavior due to early postnatal administration of morphine.

**Figure 9.**
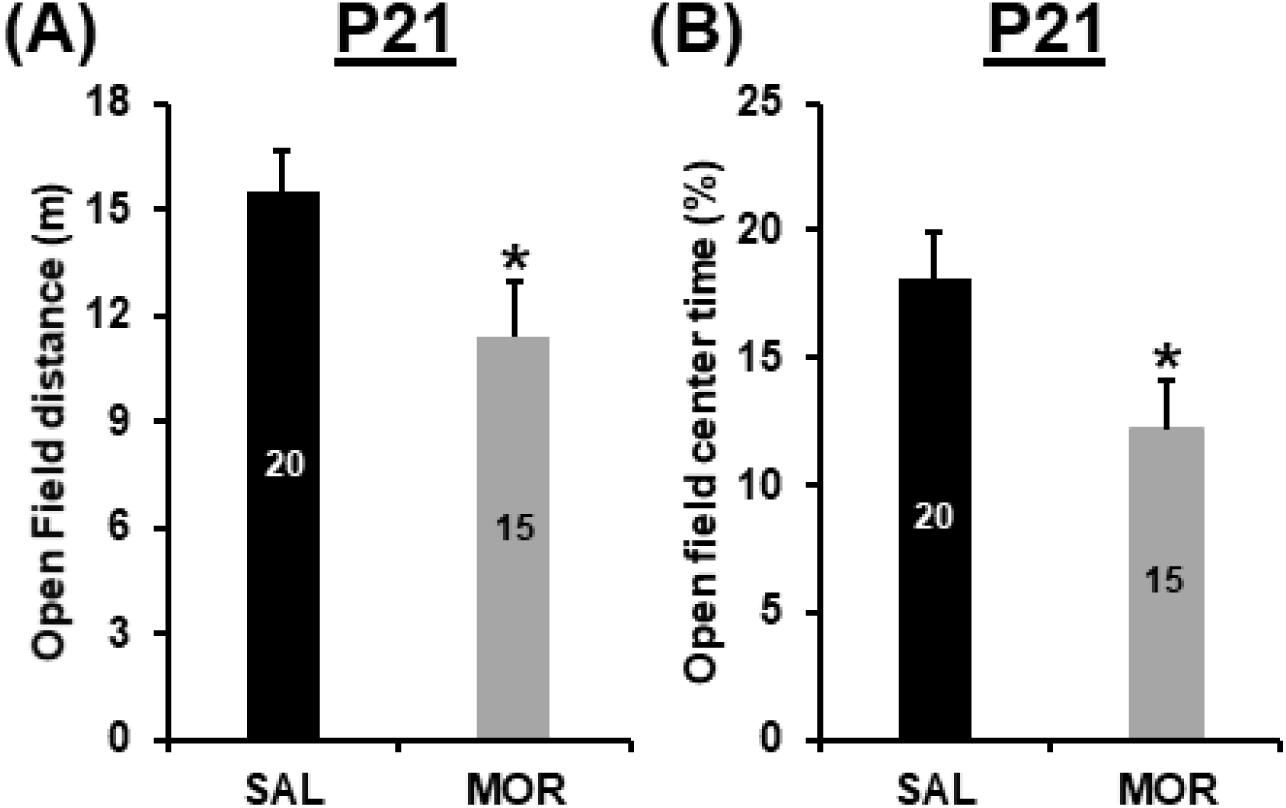
Increased anxiety-like behavior in the open field at P21 following repeated neonatal morphine exposure. **(A)** In examining distance traveled in the open field assay on P21, there was a main effect of Treatment (F_1,31_ = 4.66, p = 0.039), but not Sex (F_1,31_ = 0.07, p = 0.79) and no interaction (F_1,31_ = 0, p = 0.99). MOR mice showed less locomotor activity compared to SAL mice (t_1,33_ = 2.23, **\***p = 0.033). **(B)** In examining percent time spent in the center of the open field, the effect of Treatment just missed statistical significance (F_1,31_ = 4.15, p = 0.053). There was no effect of Sex (F_1,31_ = 0.001; p = 0.97) or Treatment x Sex interaction (F_1,31_ = 0.17; p = 0.68). Unpaired t-test indicated that MOR mice spent less percent time in the center of the open field than SAL mice (t_1,33_ = 2.10, **\***p = 0.04).

### Effect of NOWS on the brainstem transcriptome at P15

We first compared saline and morphine administration groups collapsed across females and males. We identified 123 DE genes at log_2_FC > ±0.26 and p<0.05 (Figure 10A; Extended Data Figure 10-1, 10-2). Several DE genes were of interest given their potential involvement in NOWS, including the upregulation of *Slc6a2*, which encodes the norepinephrine transporter, and downregulation of *Onecut3*, encoding for a transcription factor involved in the development of noradrenergic neurons ^42^. We investigated the biological significance of the DE genes. We identified the top enriched pathways in the morphine mice were muscle contraction, embryonic organ morphogenesis, regulation of response to food, positive regulation of gene expression, anterior/posterior pattern specification, and tissue development (adjusted p<0.05; Figure 11; Extended Data Figure 11-1).

**Figure 10.**
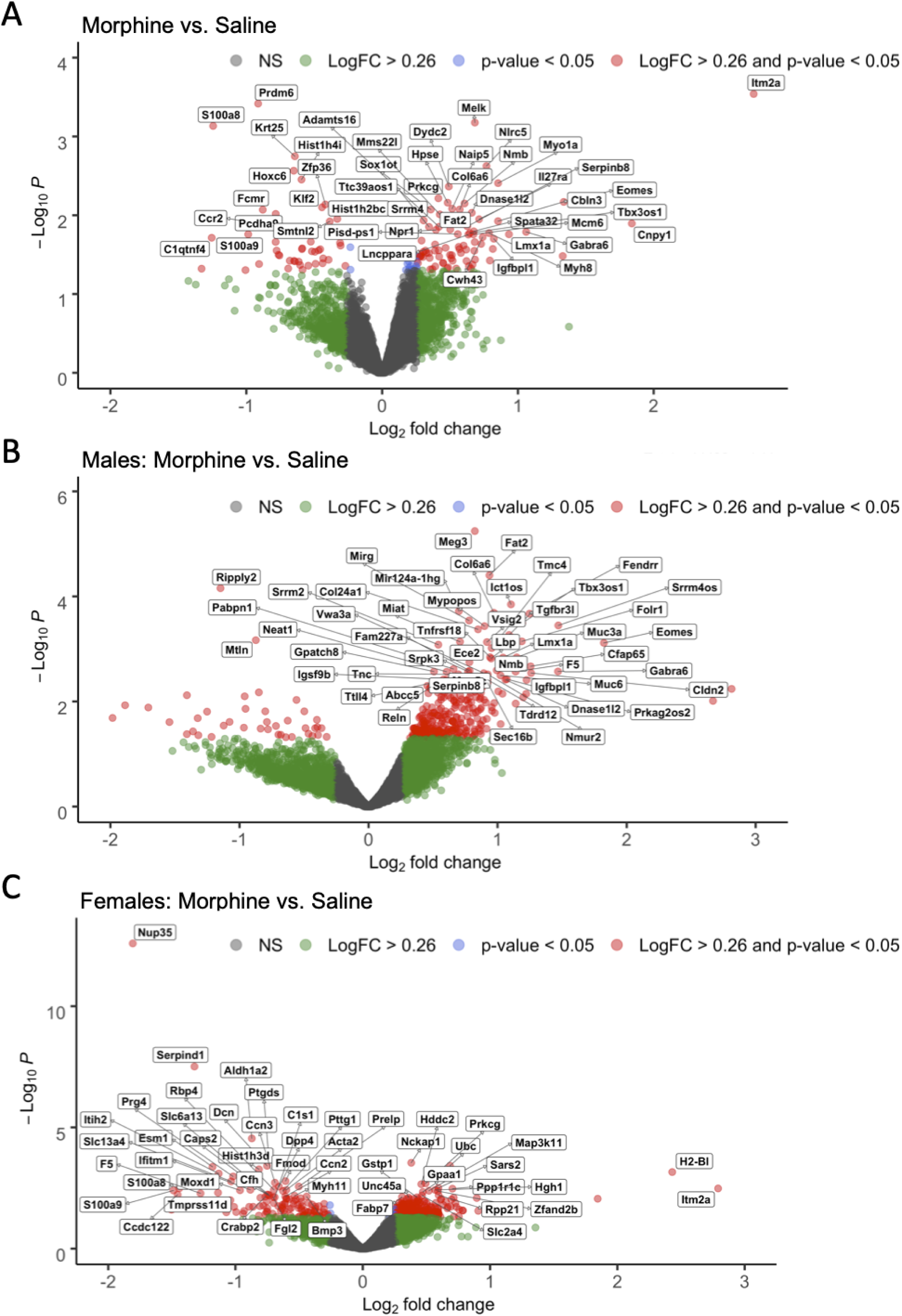
Differentially expressed genes in P15 brainstem tissue following repeated neonatal morphine exposure. The effect of morphine exposure on gene expression relative to saline mice in the sex-collapsed dataset (**A**; n=6 saline (3 male, 3 female), 6 morphine (3 male, 3 female)), males only (**B**; n=3 saline, 3 morphine), and females only (C; n=3 saline, 3 morphine). X-axis shows gene expression as Log_2_ fold change (LogFC) and y-axis shows -Log_10_ p-values (unadjusted). Color-coding of individual genes represents LogFC > ±0.26 (green), p-value < 0.05 (blue), LogFC > ±0.26 and p-value < 0.05 (red), and non-significant (NS; gray). Labeled genes represent the top 50 DE genes at LogFC > ±0.26 and p-value < 0.05.

**Figure 11.**
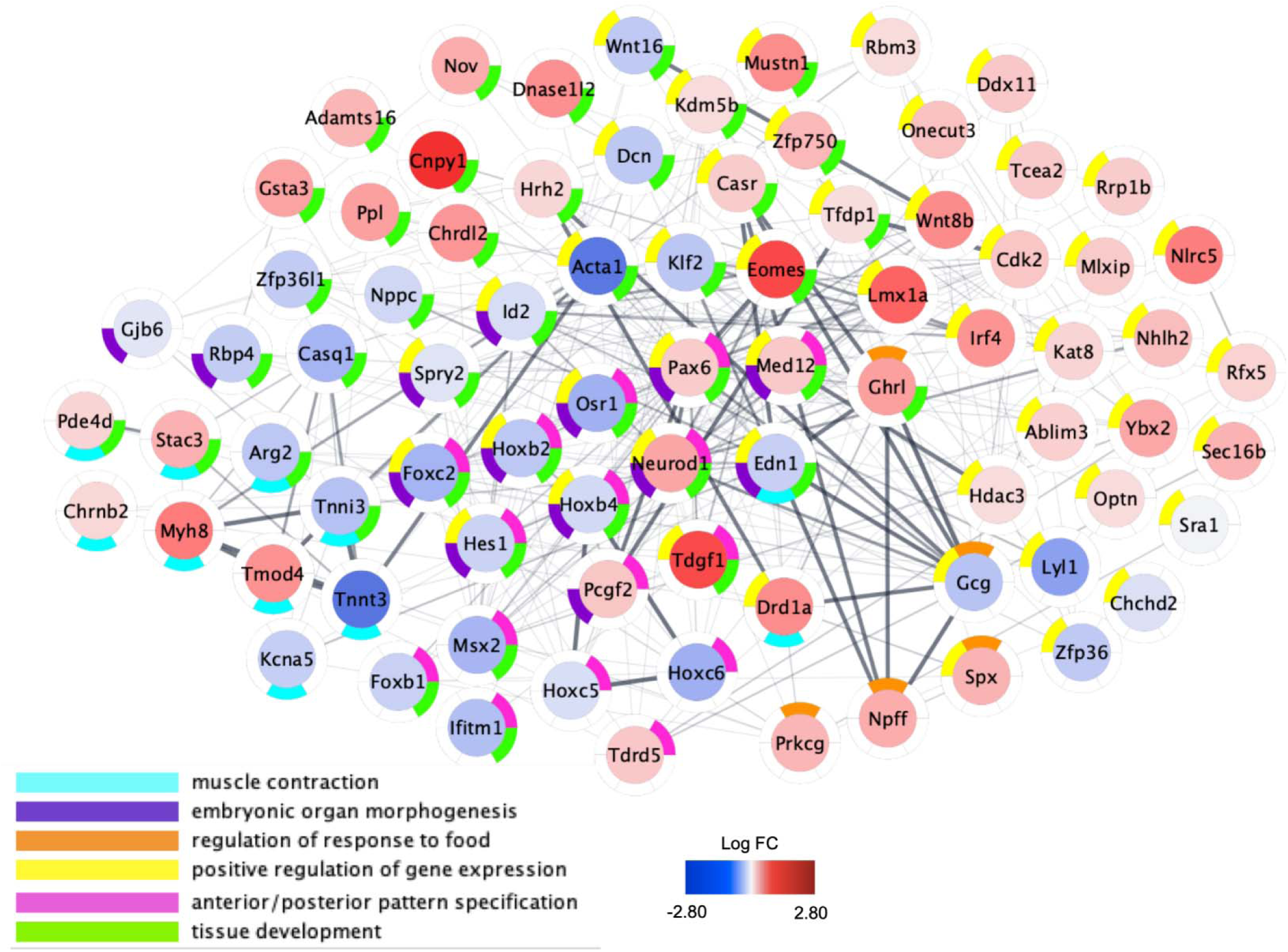
Network plot of differential brainstem gene expression within functional networks in P14 mice following repeated neonatal morphine exposure. Plots represent gene expression relative to saline mice in sex-collapsed dataset (n=6 saline (3 males, 3 females), 6 morphine (3 males, 3 females)). Central color coding of individual genes reflects Log_2_ fold change (LogFC) from -2.80 (blue) to 2.80 (red). Donut plots are color coded to reflect enriched GO Biological Process networks. All genes with unadjusted p<0.10 were included in functional network identification. Plots were generated using Cytoscape software and gene interactions were imported from the STRING database to reflect known gene interactions (shown via interconnecting lines, increased line opacity represents stronger known gene interaction).

We next explored the overall contribution of sex to the transcriptional changes in the brainstem induced by morphine administration. In females, we identified 320 DE genes at log_2_FC > ±0.26 and p<0.05 (Figure 10C; Extended Data Figure 10-3, 10-4), including the downregulation of *Nup35*, a nucleoporin gene, and *Serpind1,* part of the serpin gene superfamily that is involved in inflammation. Top enrichment terms for the females-only dataset included ribosome, oxidative phosphorylation, focal adhesion, and protein digestion and absorption, pathways in cancer, and human papillomavirus infection (Extended Data Figure 12-1). In males, we identified 328 DE genes at log_2_FC > ±0.26 and p<0.05 (Figure 10B; Extended Data Figure 10-5, 10-6), including *Meg3*, which encodes for a long noncoding RNA involved in brainstem development (Reddy et al., 2017). In addition, we found a unique set of DE genes in morphine exposed male mice that was enriched for voltage-dependent calcium channels, including the upregulation of *Cacna1c*, *Cacna1d,* and *Cacna1e*. The top enriched pathways in morphine exposed males were ribosome, oxidative phosphorylation, focal adhesion, cancer, protein digestion and absorption and human papillomavirus infection. Other pathways in males included circadian entrainment, endocannabinoid signaling, and immune function (*e.g.,* leukocyte degranulation and aggregation, B-cell malignancies, RAGE and toll-like receptor binding, Fc epsilon RI signaling, and matrix metalloproteinase-8 levels), endothelial cell function (migration, angiogenesis, VEGF signaling), and epigenetic modifications (Figure 12; Extended Data Figure 12-2).

**Figure 12:**
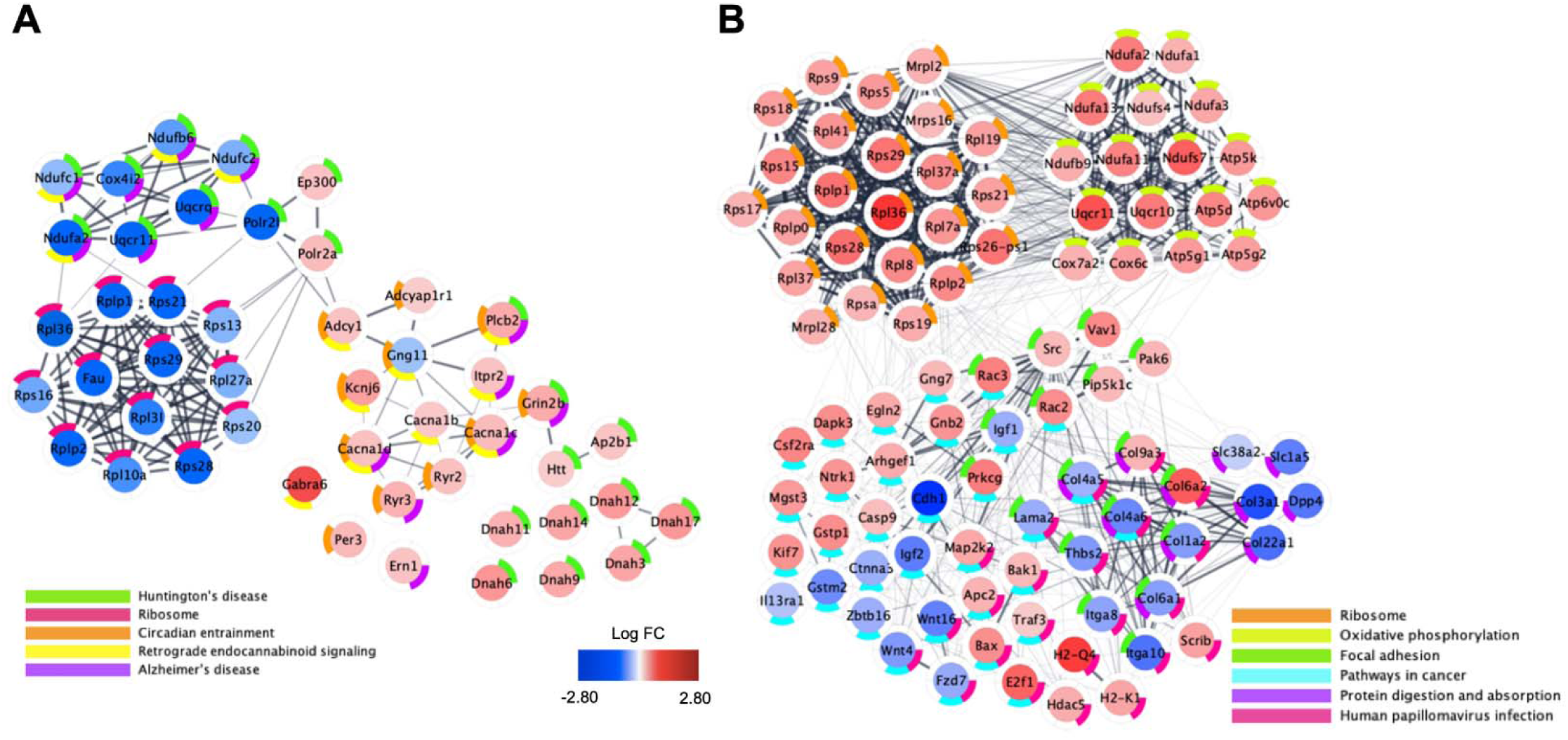
Network plots within functional networks of differential brainstem gene expression in P14 females and P14 males following repeated neonatal morphine exposure. Plots represent gene expression relative to saline exposed males (A; n=3 saline, 3 morphine) and females (B; 3 saline, 3 morphine). Central color coding of individual genes reflects Log_2_ fold change (LogFC) from -2.80 (blue) to 2.80 (red). Donut plots are color coded to reflect KEGG enrichment pathways containing fewer than 50 implicated genes. All genes with unadjusted p<0.10 were included in functional network identification. Plots were generated using Cytoscape software and gene interactions were imported from the STRING database to reflect known gene interactions (shown via interconnecting lines, increased line opacity represents stronger known gene interaction).

### NOWS impacts alternative splicing

All significant alternative splicing events based on differential intron/exon bin usage are listed in Table 2 and Table 3. There were no significant alternative splicing events for the overall effect of morphine administration. However, we identified six genes (FDR < 0.05), including *Ptov1*, *Ssbp4*, *Mmp2*, *D430019H16Rik*, *1700028K03Rik*, and *Lrch4,* that differed by sex and administration group (Table 1). Although there were no splicing events detected in the females-only dataset, several alternative splicing events were driven in part by the morphine group in males (Table 2), which included *Ptov1*, *Brdt*, *Cdkn1c*, *Tm9sf4*, *Gipc1*, *Atg2b*, *Cul9*, and *Lrch4*. In *Cdkn1c*, we detected two intronic retention events in morphine males (Table 2). *Cdkn1c* regulates the proliferation and differentiation of dopaminergic neurons in the midbrain (Andrews et al., 2007; McNamara et al., 2018). We also detected an intronic retention event within *Brdt* in males (Table 2), which codes for bromodomain testes-associated protein.

**Table 1.** Gene transcripts showing evidence for alternative splicing in the sex-combined dataset: Morphine Treatment x Sex interaction

|  |  |  |  |  |  |  |  |  |  |  |  |  |  |  |
| --- | --- | --- | --- | --- | --- | --- | --- | --- | --- | --- | --- | --- | --- | --- |
| bin_name | symbol | Gene | feature | event | locus | Locus overlap | Coordinates | start | end | length | Log FC | abs logFC | pvalue | Bin fdr |
| 84113:E038 | 84113 | Ptov1 | E | - | 84113 | - | chr7:44863067-44869788 | 44867455 | 44867512 | 58 | -2.76 | 2.76 | 2.87E-07 | 0.025 |
| 76900:E029 | 76900 | Ssbp4 | E | - | 76900 | - | chr8:70597490-70608872 | 70599525 | 70599537 | 13 | -2.93 | 2.93 | 3.96E-07 | 0.025 |
| 17390:I024 | 17390 | Mmp2 | I | - | 17390 | - | chr8:92827291-92853420 | 28546290 | 28598105 | 51816 | -1.28 | 1.28 | 4.46E-07 | 0.025 |
| 268595:I001 | 268595 | D430019H16Rik | I | - | 268595 | - | chr12:105453856-105493095 | 78291839 | 78291961 | 123 | 7.63 | 7.63 | 1.04E-06 | 0.044 |
| 76421:I007 | 76421 | 1700028K03Rik | I | - | 76421 | 231570 | chr5:107507621-107580596 | 1.33E+08 | 1.33E+08 | 484 | -8.26 | 8.26 | 1.34E-06 | 0.045 |
| 231798:E017 | 231798 | Lrch4 | E | - | 231798 | 1E+08 | chr5:137629121-137641099 | 1.38E+08 | 1.38E+08 | 2 | -4.87 | 4.87 | 1.75E-06 | 0.049 |

**Table 2.** Gene transcripts showing evidence for alternative splicing in the males-only dataset: Effect of morphine treatment

|  |  |  |  |  |  |  |  |  |  |  |  |  |
| --- | --- | --- | --- | --- | --- | --- | --- | --- | --- | --- | --- | --- |
| symbol | Gene | feature | event | locus | locus_overlap | gene_coordinates | start | end | length | logFC | pvalue | bin.fdr |
| 114642 | Brdt | I | - | 114642 | - | chr5:107331159-107387058 | 1.42E+08 | 1.42E+08 | 27158 | -1.38124 | 2.70E-08 | 0.003 |
| 12577 | Cdkn1c | I | - | 12577 | - | chr7:143458339-143461050 | 1.43E+08 | 1.43E+08 | 3842 | -1.32251 | 5.20E-08 | 0.003 |
| 12577 | Cdkn1c | I | - | 12577 | - | chr7:143458339-143461050 | 1.43E+08 | 1.43E+08 | 5304 | -1.3707 | 1.25E-07 | 0.0047 |
| 84113 | Ptov1 | E | - | 84113 | - | chr7:44863067-44869788 | 44867455 | 44867512 | 58 | 1.98688 | 1.38E-07 | 0.0047 |
| 99237 | Tm9sf4 | I | - | 99237 | - | chr2:153161303-153210466 | 73861694 | 73862277 | 584 | -2.91739 | 2.33E-07 | 0.0066 |
| 67903 | Gipc1 | I | - | 67903 | - | chr8:83652677-83664694 | 30900397 | 30900637 | 241 | -1.22149 | 1.83E-06 | 0.041 |
| 76559 | Atg2b | I | - | 76559 | - | chr12:105616136-105685211 | 30121066 | 30121475 | 410 | 3.349581 | 1.93E-06 | 0.041 |
| 78309 | Cul9 | E | - | 78309 | - | chr17:46500605-46546388 | 46521041 | 46521156 | 116 | 2.133526 | 2.59E-06 | 0.049 |
| 231798 | Lrch4 | E | - | 231798 | 1E+08 | chr5:137629121-137641099 | 1.38E+08 | 1.38E+08 | 2 | 3.464074 | 2.90E-06 | 0.0495 |

**Table 3.**
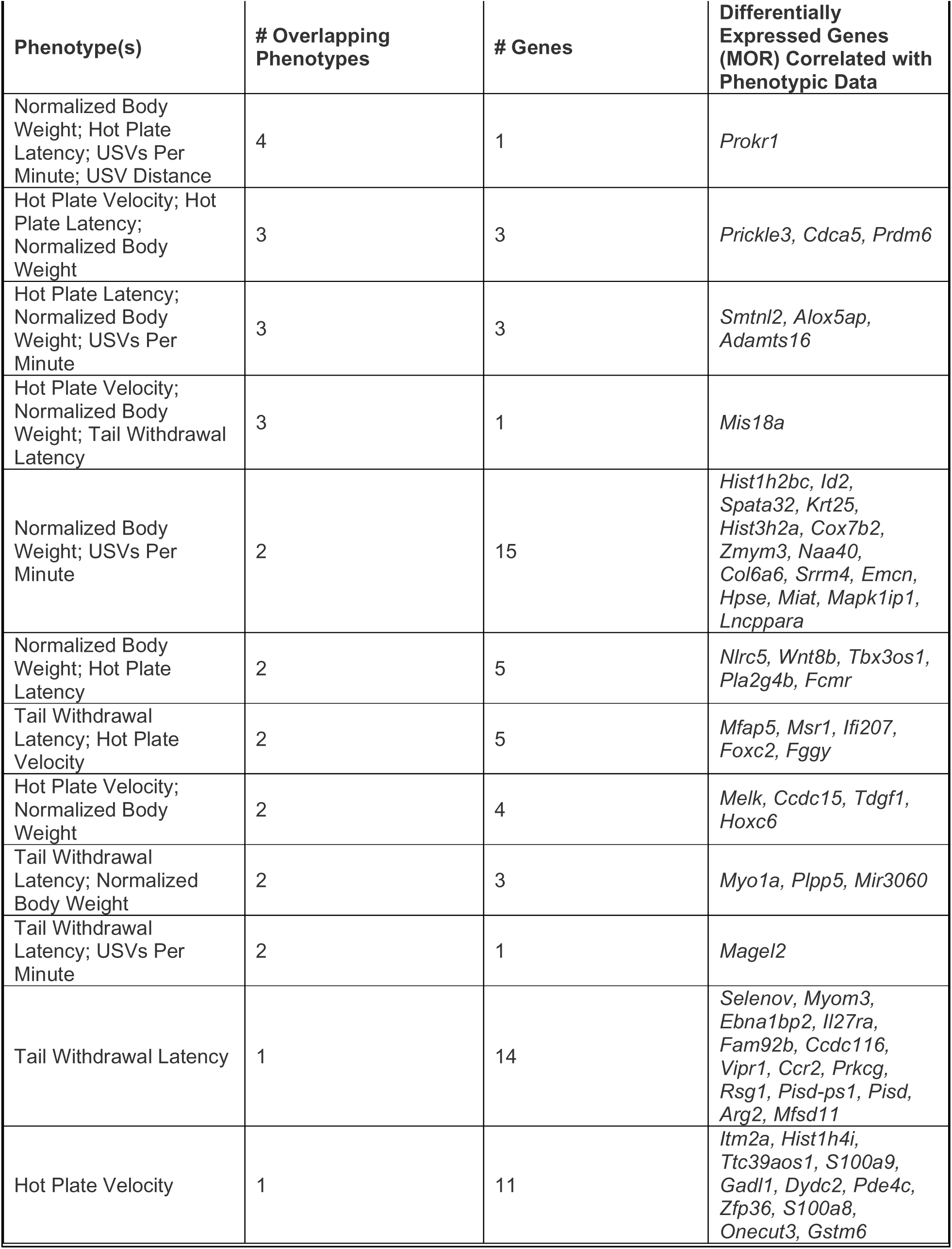
Differentially expressed gene transcripts (p < 0.05) that significantly correlated (p < 0.05) with NOWS-model phenotypes on P14

|  |  |  |  |
| --- | --- | --- | --- |
| Phenotype(s) | # Overlapping Phenotypes | # Genes | Differentially Expressed Genes (MOR) Correlated with Phenotypic Data |
| Normalized Body Weight; Hot Plate Latency; USVs Per Minute; USV Distance | 4 | 1 | *Prokr1* |
| Hot Plate Velocity; Hot Plate Latency; Normalized Body Weight | 3 | 3 | *Prickle3, Cdca5, Prdm6* |
| Hot Plate Latency; Normalized Body Weight; USVs Per Minute | 3 | 3 | *Smtnl2, Alox5ap, Adamts16* |
| Hot Plate Velocity; Normalized Body Weight; Tail Withdrawal Latency | 3 | 1 | *Mis18a* |
| Normalized Body Weight; USVs Per Minute | 2 | 15 | *Hist1h2bc, Id2, Spata32, Krt25, Hist3h2a, Cox7b2, Zmym3, Naa40, Col6a6, Srrm4, Emcn, Hpse, Miat, Mapk1ip1, Lncppara* |
| Normalized Body Weight; Hot Plate Latency | 2 | 5 | *Nlrc5, Wnt8b, Tbx3os1, Pla2g4b, Fcmr* |
| Tail Withdrawal Latency; Hot Plate Velocity | 2 | 5 | *Mfap5, Msr1, Ifi207, Foxc2, Fggy* |
| Hot Plate Velocity; Normalized Body Weight | 2 | 4 | *Melk, Ccdc15, Tdgf1, Hoxc6* |
| Tail Withdrawal Latency; Normalized Body Weight | 2 | 3 | *Myo1a, Plpp5, Mir3060* |
| Tail Withdrawal Latency; USVs Per Minute | 2 | 1 | *Magel2* |
| Tail Withdrawal Latency | 1 | 14 | *Selenov, Myom3, Ebna1bp2, Il27ra, Fam92b, Ccdc116, Vipr1, Ccr2, Prkcg, Rsg1, Pisd-ps1, Pisd, Arg2, Mfsd11* |
| Hot Plate Velocity | 1 | 11 | *Itm2a, Hist1h4i, Ttc39aos1, S100a9, Gadl1, Dydc2, Pde4c, Zfp36, S100a8, Onecut3, Gstm6* |

### DE gene correlations with NOWS behavioral phenotypes at P14

We correlated DE gene expression with behavioral phenotypes at P14. Notably, 87/139 (∼63%) of DE genes correlated with at least a single behavioral phenotype at P14 and 41/139 (∼30%) of DE genes correlated with at least two phenotypes (Table 3). Interestingly, the expression of *Prokr1* was significantly correlated with the most phenotypes, including body weight (R=-0.81, p=0.001), hot plate latency (R=-0.58, p=0.04), USVs frequency (R=0.65, p=0.02), and distance traveled during USV recording session (R=0.58, p=0.04). In morphine exposed mice, *Prokr1* was upregulated compared to saline mice, and expression was negatively correlated with hot plate latency. Thus, upregulation of *Prokr1* at P15 was associated with heightened behavioral indications of hyperalgesia at P14.

## DISCUSSION

We administered morphine from P1-P14 in mouse pups, a developmental period approximating the third trimester in human neonates, to model NOWS. We found several important NOWS-related phenotypes in female and male mice, including reduced weight gain, developmental delays in motor and USVs, agitation and increased anxiety-like behavior, and thermal hyperalgesia. We also discovered the possibility of sex-specific vulnerabilities to NOWS-model behaviors. In human neonates, males are more frequently diagnosed with and treated for NOWS, with no overall increase in NOWS severity compared to females (Juni et al., 2008). Increased male sensitivity to NOWS has also been found in C57BL/6J male mice, as evident by an increase in USVS (Robinson et al., 2019). However, our findings suggest outbred CFW female mice were more sensitive to the NOWS neurobehavioral phenotypes than males (*e.g.,* delayed latencies to self-right and increased USVs), suggesting sex-specific neurodevelopmental delays in NOWS. Motor delays and an earlier emergence of increased USVs have also been reported in C57BL/6J female mice (Robinson et al., 2019). Thus, repeated bouts of MOR intoxication and withdrawal during the third trimester-equivalent exposure is sufficient to induce multiple signs of neurodevelopmental delay. Overall, female mice were more vulnerable the effects of morphine during P1-P14 on these neurobehavioral and development phenotypes. Importantly, we also found sex-specific transcriptional alterations in the brainstem following morphine, with enrichment of mitochondrial and oxidative phosphorylation pathways enriched in females and enrichment of immune-related pathways in males. Collectively, our findings begin to highlight the consequences of morphine exposure during critical developmental periods.

Emission of USVs by pups is exhibited during distress (Hofer, 1996) and typically peak at P7 then by P14 (Elwood and Keeling, 1982). Consistent with previous studies in rats (Barr et al., 1998; Barr and Wang, 1992; Jones et al., 2002; Jones and Barr, 2001; Kalinichev and Holtzman, 2003) and mice (Robinson et al., 2019), we found an increase in USVs in morphine-versus saline-exposed mice at P14. The reinforcement of USVs during maternal separation may be mediated endogenous opioids (Hamed et al., 2012) as mu opioid receptors are required for this behavior (Moles et al., 2004). Thus, differences in endogenous opioid signaling in morphine exposed pups could contribute to the increase in USVs. Changes in USVs emerged earlier in morphine exposed female mice compared to males during opioid withdrawal. An earlier emergence of augmented USVs in female mice contrasts with prior studies using NOWS-related models where C57BL/6J inbred males were more vulnerable to consequences of morphine on USVs (Robinson et al., 2019). Inconsistencies between these studies could be due to several factors include morphine dose, frequency and interval of morphine administration, and the time course of behavioral analysis. Another factor could be use of outbred CFW mice in our studies where previous reports primarily use rats or C57BL/6J mice. Given our results, future studies using genetically diverse backgrounds of mice would be valuable for further investigating the genetic, molecular, and physiological contributions to the development of NOWS and the long-term consequences.

We also found sex-specific effects on thermal nociception in our NOWS mouse model. Previous studies reported withdrawal-induced hyperalgesia and allodynia in rat pups following early neonatal opioid exposure, including increased thermal nociception (Sweitzer et al., 2004; Zhang and Sweitzer, 2008), increased formalin-induced inflammatory nociception in P11 rats (Zissen et al., 2006) and mechanical allodynia (Sweitzer et al., 2004; Zhang and Sweitzer, 2008; Zissen et al., 2007). Here, we showed that early neonatal morphine exposure was sufficient to induce hyperalgesia. Notably, we found hyperalgesia emerged earlier in morphine exposed females. Our findings support previous work in adult outbred CD-1 female mice that showed an earlier emergence and more persistent augmentation in opioid-induced hyperalgesia (Arout et al., 2015; Holtman and Wala, 2005; Juni et al., 2010). Enhanced thermal hyperalgesia and mechanical allodynia were also observed in female rats during spontaneous opioid withdrawal, although it is unknown whether similar mechanisms in the brain lead to hyperalgesia in neonatal versus adult females. Evidence suggests that neuroadaptations of NMDA receptor signaling in males may protect against the early emergence of developmental delays. Interestingly, *Grin2b,* the gene coding for the GluN2B subunit of the NMDA receptor, was significantly upregulated only in morphine exposed male mice, consistent with male-specific morphine-induced neuroadaptations of the NMDA receptor system (Charles et al., 2017).

To investigate the potential mechanisms that contribute to these sex-specific effects of early neonatal morphine on behavior, we completed RNA-seq on the brainstems of male and female mice at P15, approximately 16 h following their final morphine injection. We identified nearly distinct sets of DE genes and enriched pathways between morphine exposed male and female mice. In females, highly significant enrichment terms were identified for 25 ribosomal genes and several genes coding for mitochondrial proteins and oxidative phosphorylation, all of which were upregulated in response to morphine. We identified the retinaldehyde dehydrogenase, *Aldh1a2*, as a significantly downregulated gene in morphine exposed females. ALDH1A2 synthesizes retinoic acid during early stages of development and has been linked to neural tube defects (Niederreither et al., 2002). Another gene involved in neurogenesis and cortical development, *Cdh1* (Delgado-Esteban et al., 2013), was also reduced in the brainstems of morphine-exposed female mice, suggesting morphine exposure and spontaneous withdrawal from morphine may disrupt neurodevelopmental processes by altering the expression of key genes. In addition, we identified upregulation of genes involved in nociception, including *Pnoc*, a gene coding for the precursor protein to nociception. Nociceptin levels in the spinal cord were higher in female rats treated with morphine (Zhang et al., 2012). Together, our findings support the notion that early postnatal exposure to morphine leads to changes in nociceptin signaling, although the potential role in hyperalgesia during opioid withdrawal in neonates is unknown.

Enrichment of pathways in male mice were largely distinct from those of females exposed to morphine. We observed several pathways enriched in male mice following morphine exposure, including circadian entrainment. Within the circadian entrainment pathway, the following genes were significantly upregulated in morphine exposed male mice: *Per3* (circadian gene); *Adcyap1r1* (PACAP1 receptor); *Ryr2,3* (intracellular calcium channels); *Cacna1c,d,e* (membrane voltage-gated calcium channels); *Grin2b* (NMDA receptor subunit); *Kcnj6* (Kir3.2/GIRK2 potassium channel); and *Kcnj13* (Kir7.1 potassium channel). Chronic morphine altered the expression of calcium and potassium channels in the mouse cortex and limbic system that contributes to opioid withdrawal, tolerance and hyperalgesia(Cruz et al., 2008; Lee et al., 2011; Shibasaki et al., 2011, 2010, 2007). Importantly, circadian genes, such as *Per3*, which was changed in morphine-exposed male mice, have been associated with opioid-induced hyperalgesia in mice (Zhang et al., 2019) and mutations in *PER3* are associated with opioid dependence in humans (Surovtseva et al., 2012). Other genes that were upregulated in morphine exposed males include the PACAP1 receptor and ryanodine receptors, which are also associated with opioid withdrawal (Lipták et al., 2012; Mácsai et al., 2002; Martin et al., 2003)(Ohsawa and Kamei, 1999).

In both males and females, we found an upregulation of *Slc6a2*, which encodes for the norepinephrine transporter (NET). This transcriptional adaptation might serve to counter the increase in extracellular norepinephrine levels, a hallmark feature of opioid withdrawal (Koob, 2019). Indeed, it is well-established that reducing norepinephrine release by pharmacological agonism at the alpha-2 receptor with clonidine can alleviate the severity of the sympathetic component of opioid withdrawal (Gowing et al., 2016). Our findings suggest that clonidine might also be an effective treatment for autonomic symptoms in NOWS.

Human neonates born with NOWS often have a lower birth weight and have difficulty with post-partum weight gain. Consistent with other mouse models of NOWS (Robinson et al., 2019), we found a significant reduction in weight gain in morphine-exposed pups that persisted from P14 through weaning into adulthood. Reduced weight gain can be caused by several factors including the compromised ability to feed or reduced motivation to feed. Opioids inhibit gastrointestinal motility, inducing nausea and leading to anorexia (Santolaria-Fernández et al., 1995). Opioid-induced weight loss exerts profound effects on nutrition, health, and physiology in OUD (Santolaria-Fernández et al., 1995) and may be a hallmark in the transition to opioid dependence (Chen et al., 2006).

Collectively, we demonstrated sex-specific phenotypes using a NOWS model in outbred mice and revealed potential brainstem mechanisms of opioid withdrawal severity related to neurodevelopmental delays. Our studies serve as the foundation for future endeavors to examine the transcriptional changes in additional brain regions and specific cell types related to NOWS-related behavioral phenotypes. Our work establishes divergent neurobiological adaptations between males and females that could have implications for treatment and prevention of NOWS. Finally, because the sex differences we have identified contrast with previous reports on different genetic backgrounds (Robinson et al., 2019), they suggest a potential genetic component that should be further explored on multiple genetic backgrounds.

## Supporting information

Differentially expressed genes, sex-combined dataset

Differentially expressed genes, sex-combined dataset (log2FC>0.26; p<0.05)

Differentially expressed genes, females-only dataset

Differentially expressed genes, females-only dataset (log2FC>0.26; p<0.05)

Differentially expressed genes, males-only dataset

Differentially expressed genes, males-only dataset (log2FC>0.26; p<0.05)

Enrichment analysis (sex-combined)

Enrichment analysis (females)

Enrichment analysis (males)

## ACKNOWLEDGEMENTS

Spivack Center for Clinical and Translational Neuroscience (C.D.B.), NIH/NIDA R01DA039168 (C.D.B.), NIH/NIDA U01DA050243 (C.D.B.), NIH/NICHD R01HD096798 (E.W.), Brain and Behavior Research Foundation (A.C.M.), Boston University’s Undergraduate Research Opportunities Program (J.L.S.), and Dr. Pieter Faber at the University of Chicago Genomics Facility

## REFERENCES

Andrews SC, Wood MD, Tunster SJ, Barton SC, Surani MA, John RM (2007) Cdkn1c (p57Kip2) is the major regulator of embryonic growth within its imprinted domain on mouse distal chromosome 7. BMC Dev Biol 7:53.

Arout CA, Caldwell M, Rossi G, Kest B (2015) Spinal and supraspinal N-methyl-D-aspartate and melanocortin-1 receptors contribute to a qualitative sex difference in morphine-induced hyperalgesia. Physiol Behav 147:364–372.

Barr GA, McPhie-Lalmansingh A, Perez J, Riley M (2011) Changing mechanisms of opiate tolerance and withdrawal during early development: animal models of the human experience. ILAR J 52:329–341.

Barr GA, Wang S (1992) Tolerance and withdrawal to chronic morphine treatment in the week-old rat pup. Eur J Pharmacol 215:35–42.

Barr GA, Zmitrovich A, Hamowy AS, Liu PY, Wang S, Hutchings DE (1998) Neonatal withdrawal following pre- and postnatal exposure to methadone in the rat. Pharmacol Biochem Behav 60:97–104.

Bederson JB, Fields HL, Barbaro NM (1990) Hyperalgesia during naloxone-precipitated withdrawal from morphine is associated with increased on-cell activity in the rostral ventromedial medulla. Somatosens Mot Res 7:185–203.

Bertrand A, Demuynck K, Stouten V (2008) UNSUPERVISED LEARNING OF AUDITORY FILTER BANKS USING NON-NEGATIVE MATRIX FACTORISATION. International Conference on Acoustics, Speech, and Signal Processing 4713– 4716.

Boasen JF, McPherson RJ, Hays SL, Juul SE, Gleason CA (2009) Neonatal stress or morphine treatment alters adult mouse conditioned place preference. Neonatology 95:230–239.

Bolger AM, Lohse M, Usadel B (2014) Trimmomatic: a flexible trimmer for Illumina sequence data. Bioinformatics 30:2114–2120.

Byrnes EM, Vassoler FM (2018) Modeling prenatal opioid exposure in animals: Current findings and future directions. Front Neuroendocrinol 51:1–13.

Charles MK, Cooper WO, Jansson LM, Dudley J, Slaughter JC, Patrick SW (2017) Male Sex Associated With Increased Risk of Neonatal Abstinence Syndrome. Hosp Pediatr 7:328–334.

Chen H-H, Chiang Y-C, Yuan ZF, Kuo C-C, Lai M-D, Hung T-W, Ho I-K, Chen S-T (2015) Buprenorphine, methadone, and morphine treatment during pregnancy: behavioral effects on the offspring in rats. Neuropsychiatr Dis Treat 11:609–618.

Chen SA, O’Dell LE, Hoefer ME, Greenwell TN, Zorrilla EP, Koob GF (2006) Unlimited access to heroin self-administration: independent motivational markers of opiate dependence. Neuropsychopharmacology 31:2692–2707.

Cruz HG, Berton F, Sollini M, Blanchet C, Pravetoni M, Wickman K, Lüscher C (2008) Absence and rescue of morphine withdrawal in GIRK/Kir3 knock-out mice. J Neurosci 28:4069–4077.

Delgado-Esteban M, García-Higuera I, Maestre C, Moreno S, Almeida A (2013) APC/C-Cdh1 coordinates neurogenesis and cortical size during development. Nat Commun 4:2879.

Desai RJ, Huybrechts KF, Hernandez-Diaz S, Mogun H, Patorno E, Kaltenbach K, Kerzner LS, Bateman BT (2015) Exposure to prescription opioid analgesics in utero and risk of neonatal abstinence syndrome: population based cohort study. BMJ 350:h2102.

Dobin A, Davis CA, Schlesinger F, Drenkow J, Zaleski C, Jha S, Batut P, Chaisson M, Gingeras TR (2013) STAR: ultrafast universal RNA-seq aligner. Bioinformatics 29:15–21.

Elwood RW, Keeling F (1982) Temporal organization of ultrasonic vocalizations in infant mice. Dev Psychobiol 15:221– 227.

Fodor A, Tímár J, Zelena D (2014) Behavioral effects of perinatal opioid exposure. Life Sci 104:1–8.

Gonzales NM, Palmer AA (2014) Fine-mapping QTLs in advanced intercross lines and other outbred populations. MammGenome 25:271–292.

Gowing L, Farrell M, Ali R, White JM (2016) Alpha₂-adrenergic agonists for the management of opioid withdrawal. Cochrane Database Syst Rev CD002024.

Hamed A, Taracha E, Szyndler J, Krząścik P, Lehner M, Maciejak P, Skórzewska A, Płaźnik A (2012) The effects of morphine and morphine conditioned context on 50 kHz ultrasonic vocalisation in rats. Behav Brain Res 229:447– 450.

Hofer MA (1996) Multiple regulators of ultrasonic vocalization in the infant rat. Psychoneuroendocrinology 21:203–217.

Holtman JR, Wala EP (2005) Characterization of morphine-induced hyperalgesia in male and female rats. Pain 114:62–70.

Jansson LM, Patrick SW (2019) Neonatal Abstinence Syndrome. Pediatr Clin North Am 66:353–367.

Joder C, Schuller B (2012) Exploring Nonnegative Matrix Factorization for Audio Classification: Application to Speaker Recognition. Speech Communication; ITG Symposium; Proceedings of 1–4.

Jones KL, Barr GA (2001) Injections of an opioid antagonist into the locus coeruleus and periaqueductal gray but not the amygdala precipitates morphine withdrawal in the 7-day-old rat. Synapse 39:139–151.

Jones KL, Barr GA (1995) Ontogeny of morphine withdrawal in the rat. Behav Neurosci 109:1189–1198.

Jones KL, Zhu H, Jenab S, Du T, Inturrisi CE, Barr GA (2002) Attenuation of acute morphine withdrawal in the neonatal rat by the competitive NMDA receptor antagonist LY235959. Neuropsychopharmacology 26:301–310.

Juni A, Cai M, Stankova M, Waxman AR, Arout C, Klein G, Dahan A, Hruby VJ, Mogil JS, Kest B (2010) Sex-specific mediation of opioid-induced hyperalgesia by the melanocortin-1 receptor. Anesthesiology 112:181–188.

Juni A, Klein G, Kowalczyk B, Ragnauth A, Kest B (2008) Sex differences in hyperalgesia during morphine infusion: effect of gonadectomy and estrogen treatment. Neuropharmacology 54:1264–1270.

Kalinichev M, Holtzman SG (2003) Changes in urination/defecation, auditory startle response, and startle-induced ultrasonic vocalizations in rats undergoing morphine withdrawal: similarities and differences between acute and chronic dependence. J Pharmacol Exp Ther 304:603–609.

Kaplan H, Fields HL (1991) Hyperalgesia during acute opioid abstinence: evidence for a nociceptive facilitating function of the rostral ventromedial medulla. J Neurosci 11:1433–1439.

Koob GF (2019) Neurobiology of Opioid Addiction: Opponent Process, Hyperkatifeia, and Negative Reinforcement. Biol Psychiatry.

Lee M, Silverman SM, Hansen H, Patel VB, Manchikanti L (2011) A comprehensive review of opioid-induced hyperalgesia. Pain Physician 14:145–161.

Liao Y, Smyth GK, Shi W (2019) The R package Rsubread is easier, faster, cheaper and better for alignment and quantification of RNA sequencing reads. Nucleic Acids Res 47:e47.

Lipták N, Dochnal R, Babits A, Csabafi K, Szakács J, Tóth G, Szabó G (2012) The effect of pituitary adenylate cyclase-activating polypeptide on elevated plus maze behavior and hypothermia induced by morphine withdrawal. Neuropeptides 46:11–17.

Mácsai M, Pataki I, Tóth G, Szabó G (2002) The effects of pituitary adenylate cyclase-activating polypeptide on acute and chronic morphine actions in mice. Regul Pept 109:57–62.

Maeda T, Kishioka S, Inoue N, Shimizu N, Fukazawa Y, Ozaki M, Yamamoto H (2002) Naloxone-precipitated morphine withdrawal elicits increases in c-fos mRNA expression in restricted regions of the infant rat brain. Jpn J Pharmacol 90:270–275.

Mancini E, Rabinovich A, Iserte J, Yanovsky M, Chernomoretz A (2020) ASpli: Analysis of alternative splicing using RNA-Seq.

Martin M, Otto C, Santamarta MT, Torrecilla M, Pineda J, Schütz G, Maldonado R (2003) Morphine withdrawal is modified in pituitary adenylate cyclase-activating polypeptide type I-receptor-deficient mice. Brain Res Mol Brain Res 110:109–118.

McNamara GI, Davis BA, Browne M, Humby T, Dalley JW, Xia J, John RM, Isles AR (2018) Dopaminergic and behavioural changes in a loss-of-imprinting model of Cdkn1c. Genes Brain Behav 17:149–157.

McPhie AA, Barr GA (2009) Regional Fos expression induced by morphine withdrawal in the 7-day-old rat. Dev Psychobiol 51:544–552.

Milliren CE, Gupta M, Graham DA, Melvin P, Jorina M, Ozonoff A (2018) Hospital Variation in Neonatal Abstinence Syndrome Incidence, Treatment Modalities, Resource Use, and Costs Across Pediatric Hospitals in the United States, 2013 to 2016. Hosp Pediatr 8:15–20.

Minakova E, Sarafinovska S, Mikati MO, Barclay KM, McCullough KB, Dougherty JD, Al-Hasani R, Maloney SE (2021) Ontogenetic Oxycodone Exposure Affects Early Life Communicative Behaviors, Sensorimotor Reflexes, and Weight Trajectory in Mice. Front Behav Neurosci 15:615798.

Moles A, Kieffer BL, D’Amato FR (2004) Deficit in attachment behavior in mice lacking the mu-opioid receptor gene. Science 304:1983–1986.

Niederreither K, Abu-Abed S, Schuhbaur B, Petkovich M, Chambon P, Dollé P (2002) Genetic evidence that oxidative derivatives of retinoic acid are not involved in retinoid signaling during mouse development. Nat Genet 31:84– 88.

Noirot E (1966) Ultra-sounds in young rodents. I. Changes with age in albino mice. Anim Behav 14:459–462.

Ohsawa M, Kamei J (1999) Role of intracellular calcium on the modulation of naloxone-precipitated withdrawal jumping in morphine-dependent mice by diabetes. Brain Res 815:424–430.

Ossipov MH, Lai J, King T, Vanderah TW, Porreca F (2005) Underlying mechanisms of pronociceptive consequences of prolonged morphine exposure. Biopolymers 80:319–324.

Parker CC, Gopalakrishnan S, Carbonetto P, Gonzales NM, Leung E, Park YJ, Aryee E, Davis J, Blizard DA, Ackert-Bicknell CL, Lionikas A, Pritchard JK, Palmer AA (2016) Genome-wide association study of behavioral, physiological and gene expression traits in outbred CFW mice. NatGenet.

Reddy AS, O’Brien D, Pisat N, Weichselbaum CT, Sakers K, Lisci M, Dalal JS, Dougherty JD (2017) A Comprehensive Analysis of Cell Type-Specific Nuclear RNA From Neurons and Glia of the Brain. Biol Psychiatry 81:252–264.

Robinson MD, McCarthy DJ, Smyth GK (2010) edgeR: a Bioconductor package for differential expression analysis of digital gene expression data. Bioinformatics 26:139–140.

Robinson SA, Jones AD, Brynildsen JK, Ehrlich ME, Blendy JA (2019) Neurobehavioral effects of neonatal opioid exposure in mice: Influence of the OPRM1 SNP. Addict Biol e12806.

Ryan G, Dooley J, Gerber Finn L, Kelly L (2019) Nonpharmacological management of neonatal abstinence syndrome: a review of the literature. J Matern Fetal Neonatal Med 32:1735–1740.

Santolaria-Fernández FJ, Gómez-Sirvent JL, González-Reimers CE, Batista-López JN, Jorge-Hernández JA, Rodríguez-Moreno F, Martínez-Riera A, Hernández-García MT (1995) Nutritional assessment of drug addicts. Drug Alcohol Depend 38:11–18.

Semple BD, Blomgren K, Gimlin K, Ferriero DM, Noble-Haeusslein LJ (2013) Brain development in rodents and humans: Identifying benchmarks of maturation and vulnerability to injury across species. Prog Neurobiol 106–107:1–16.

Shibasaki M, Katsura M, Kurokawa K, Torigoe F, Ohkuma S (2007) Regional differences of L-type high voltage-gated calcium channel subunit expression in the mouse brain after chronic morphine treatment. J Pharmacol Sci 105:177–183.

Shibasaki M, Kurokawa K, Mizuno K, Ohkuma S (2011) Up-regulation of Ca(v)1.2 subunit via facilitating trafficking induced by Vps34 on morphine-induced place preference in mice. Eur J Pharmacol 651:137–145.

Shibasaki M, Kurokawa K, Ohkuma S (2010) Upregulation of L-type Ca(v)1 channels in the development of psychological dependence. Synapse 64:440–444.

Surovtseva MN, Kudryavtseva EA, Voronina EN, Pronin SV, Filipenko ML (2012) Association of the 54-nucleotide repeat polymorphism of hPer3 with opioid dependence in residents of the West Siberian region. Psychiatr Genet 22:309–310.

Sutter MB, Leeman L, Hsi A (2014) Neonatal opioid withdrawal syndrome. Obstet Gynecol Clin North Am 41:317–334.

Sweitzer SM, Allen CP, Zissen MH, Kendig JJ (2004) Mechanical allodynia and thermal hyperalgesia upon acute opioid withdrawal in the neonatal rat. Pain 110:269–280.

Tuttle AH, Philip VM, Chesler EJ, Mogil JS (2018) Comparing phenotypic variation between inbred and outbred mice. Nat Methods 15:994–996.

Van Segbroeck M, Knoll AT, Levitt P, Narayanan S (2017) MUPET-Mouse Ultrasonic Profile ExTraction: A Signal Processing Tool for Rapid and Unsupervised Analysis of Ultrasonic Vocalizations. Neuron 94:465–485.e5.

Vanderah TW, Suenaga NM, Ossipov MH, Malan TP, Lai J, Porreca F (2001) Tonic descending facilitation from the rostral ventromedial medulla mediates opioid-induced abnormal pain and antinociceptive tolerance. J Neurosci 21:279–286.

Vera-Portocarrero LP, Ossipov MH, Lai J, King T, Porreca F (2011) Descending facilitatory pathways from the rostroventromedial medulla mediate naloxone-precipitated withdrawal in morphine-dependent rats. J Pain 12:667–676.

Wachman EM, Schiff DM, Silverstein M (2018) Neonatal Abstinence Syndrome: Advances in Diagnosis and Treatment. JAMA 319:1362–1374.

Wang Z, Hu J, Johnson WE, Campbell JD (2019) scruff: an R/Bioconductor package for preprocessing single-cell RNA-sequencing data. BMC Bioinformatics 20:222.

Xie JY, Herman DS, Stiller C-O, Gardell LR, Ossipov MH, Lai J, Porreca F, Vanderah TW (2005) Cholecystokinin in the rostral ventromedial medulla mediates opioid-induced hyperalgesia and antinociceptive tolerance. J Neurosci 25:409–416.

Yazdani N, Parker CC, Shen Y, Reed ER, Guido MA, Kole LA, Kirkpatrick SL, Lim JE, Sokoloff G, Cheng R, Johnson WE, Palmer AA, Bryant CD (2015) Hnrnph1 Is A Quantitative Trait Gene for Methamphetamine Sensitivity. PLoS Genet 11:e1005713.

Zhang GH, Sweitzer SM (2008) Neonatal morphine enhances nociception and decreases analgesia in young rats. Brain Res 1199:82–90.

Zhang P, Moye LS, Southey BR, Dripps I, Sweedler JV, Pradhan A, Rodriguez-Zas SL (2019) Opioid-Induced Hyperalgesia Is Associated with Dysregulation of Circadian Rhythm and Adaptive Immune Pathways in the Mouse Trigeminal Ganglia and Nucleus Accumbens. Mol Neurobiol.

Zhang Y, Donica CL, Standifer KM (2012) Sex differences in the Nociceptin/Orphanin FQ system in rat spinal cord following chronic morphine treatment. Neuropharmacology 63:427–433.

Zissen MH, Zhang G, Kendig JJ, Sweitzer SM (2006) Acute and chronic morphine alters formalin pain in neonatal rats. Neurosci Lett 400:154–157.

Zissen MH, Zhang G, McKelvy A, Propst JT, Kendig JJ, Sweitzer SM (2007) Tolerance, opioid-induced allodynia and withdrawal associated allodynia in infant and young rats. Neuroscience 144:247–262.

